# Comparing Deep Learning Models for Age Prediction Based on the Resting State fMRI Dataset from the “Brain Dock” Service in Japan

**DOI:** 10.1101/2022.08.14.503923

**Authors:** Yutaka Minowa, Keiichi Onoda, Masahiro Takamura, Shuhei Yamaguchi, Naoki Okamoto, Hiroyuki Akama

## Abstract

In Japan, many hospitals provide the unequaled service of medical check-up called the “Brain Dock”; however, there is a paucity of studies aimed at leveraging functional magnetic resonance imaging (fMRI) in hospitals. We obtained the resting-state fMRI (rs-fMRI) scans of about 695 patients who accessed this service for preventive medicine against Alzheimer’s Disease. In this study, we created deep learning models for age prediction with the ultimate aim of arriving at standard protocols for introducing rs-fMRI into clinical settings, particularly in Brain Docks. With that view, an assemblage of modeling conditions was attempted, changing multiple parameter values, features based on data extraction methods (region of interest-wise mean blood-oxygen-level-dependent lump-sum time series models or dynamic functional connectivity models), deep learning algorithms (Transformer, Multi-task Transformer, and unidirectional or bidirectional long short-term memory models), and different atlas-dependent brain region segmentation methods (including the Automated Anatomical Labeling and Harvard-Oxford atlases). As a result, a robust and highly significant correlation was obtained between actual and predicted ages from all types of methodologies. In addition, we determined that some conditions had a relatively large impact on prediction performance based on extended comparisons. The accuracy decreased, particularly according to the choice of atlases but with the same modeling conditions. Notwithstanding, we found that atlases based on intrinsic functional connectivity provided significant prediction accuracy even with a small number of regions to a similar extent as networks lowered spatial granularity. Moreover, we found that multi-task learning with other phenotype data (related to gender differences) was possible, but did not improve the prediction accuracy as much as expected. Despite these limitations, our results could provide a hopeful prospect of introducing fMRI into the field of neuro-clinical practice.

**Highlights:**

1. Age prediction modeling was performed using the rs-fMRI data of a Japanese “Brain Dock” service targeting mainly elderly people, which made prediction more challenging.
2. A robust and highly significant correlation was obtained from all types of methodologies changing data extraction methods, deep learning algorithms, and brain atlases.
3. Age prediction was successful even with a very small number of regions exclusively based on intrinsic functional connectivity networks, although there was a difference in the achievement of modeling.

## 1. Introduction

The global population of dementia patients has been predicted to exceed 81 million by 2040, with Alzheimer’s disease being the most common cause (Chu, 2012; Voce et al., 2012). One way to address these problems of aging, which have significant social and economic consequences, is to take advantage of neuroimaging techniques to regularly track changes in the individual brain and detect neural diseases from an early stage. It has been suggested that machine learning models would be useful in preventive medicine in that they would allow the discernment and prediction of the preclinical and clinical stages of Alzheimer’s disease (Sarraf & Tofighi, 2016). Brain age prediction has potential neuroscientific and clinical value, and age prediction using deep learning is considered an effective tool for assessing brain health among individuals (Cole & Franke, 2017).

Japan has a unique medical checkup system called “Brain Dock (Brain Screening)” (Morita, 2019). This system is intended to “detect asymptomatic neural or cerebrovascular diseases or their risk factors, and prevent their onset or progression, mainly through imaging examinations using magnetic resonance imaging (MRI) and magnetic resonance angiography for asymptomatic subjects” (Brain Dock Guidelines, 2008). The target population of this program is mainly healthy elderly people, including some middle-aged people, for whom it serves as a regular health checkup system. A large study of older adults has suggested a significant relationship between brain age-predicted results and the risk of mortality (Cole et al., 2017a), drawing attention to the possibility that predicted brain age can be used as a biomarker of aging (Cole et al. al., 2017b).

In the present study, data from the Brain Dock of a total of 695 patients at Shimane Institute of Health Science was used to compute a model for predicting brain age. These data are unique in that they included resting-state functional MRI (fMRI) data in the scan protocol, which set it specifically apart from the other general data contained in the Brain Dock. In addition, many previous studies of age prediction have used datasets centered on younger age groups or have included younger age groups in the experimentally designed sample (Dosenbach et al., 2010; Qin et al., 2015; Li et al., 2018; Niu et al., 2019; Gadgil et al., 2020). However, the Brain Dock dataset used in this study was centered on older age groups based on the necessity of clinical practice.

From the methodological point of view mentioned above, deep learning models hold great promise for age prediction (Cole et al, 2017b). In addition to representing the hidden layers of a neural network in a data-driven manner, it also has the potential to more accurately identify previously unknown relationships. Further improvements in accuracy can be anticipated for brain age prediction. A recent study has successfully improved age prediction by applying multimodal brain imaging data to data on cortical anatomy and whole-brain functional connectivity (Liem et al., 2017). However, there is room for improvement within the realms of possibility for each modal analysis of MRI data. In particular, resting-state functional connectivity (rsFC) analysis would be fruitful if enhanced by recurrent neural networks (RNNs) such as the long short-term memory (LSTM) and Transformer neural networks, although the number of studies remains small and research in this regard is currently just getting started (Malkiel et al., 2021). Also, to the best of our knowledge, no previous studies have examined the robustness of age prediction for the same rsFC data with multiple parameter values, features, deep learning algorithms, and different atlas-dependent brain region segmentation methods. In particular, no previous reports have compared the results of models dealing with inputs from two data sources: the whole-time series of blood-oxygen-level-dependent (BOLD) signal change in each (region of interest [ROI]) on one hand, and its dynamic modulation across each of the segmented periods by sliding one time window on the other. Moreover, this is the first comprehensive study of rsFC-based age prediction, aimed at the prospect of a standard protocol that will be necessary for the introduction of fMRI into clinical settings, particularly for Brain Docks. Furthermore, the present study introduced multi-task learning (Ruder, 2017; Huang et al., 2020) for age prediction, assuming the effects of reducing the risk of overlearning as well as seeking the adaptability of the models to random noise.

## 2. Materials and methods

### 2.1. MRI: Participants, data acquisition, and preprocessing

This study utilized the data from Brain Docks for a total of 695 patients at Shimane University School of Medicine. The study protocol was first approved by the ethics committee of the Faculty of Medicine at Shimane University, and subsequently by the institutional review boards of all the organizations to which the authors are affiliated. We analyzed the data of 615 subjects, excluding the participants of experiments who had undergone multiple fMRI imaging sessions and those who were extreme outliers from the age distribution. The 615 (333 males and 282 females) consisted of 37 subjects in their 30s, 97 in their 40s, 107 in their 50s, 156 in their 60s, 166 in their 70s, and 52 in their 80s. The mean age was 62.4±13.5 years, which implied that the dataset focused on older subjects. Furthermore, the dataset was rich in test scores on neural and cognitive decline prevalent in elderly people, including mini-mental state scores, Kohs (cube combination test) values, self-rated depressive scale scores, motivation scores (on the apathy scale), diagnoses of mental disorders (according to the Diagnostic and Statistical Manual of Mental Disorders of the American Psychiatric Association), and Frontal Assessment Battery frontal lobe function test scores.

The following scan parameters for rs-fMRI images of the brain were acquired from the Philips 3.0T machine installed at Shimane Institute of Health Science in 2016: TR=2500 ms, TE=30 ms, FOV with (RL=212 mm, AP=212 mm, FH=159.2 mm), ACQ matrix M×P =64×63, slices=40, and slice thickness=3.2 mm. The rs-fMRI scan time for each participant was 350 seconds; thus, each session yielded 140 pieces of 3D functional images. The parameter values set for the structural scans using MP RAGE were as follows; TR=6.8 ms, TE=3.1 ms, flip angle=9°, bandwidth=289 Hz/Px, and voxel size= 1×1×1.2 mm. The DICOM data were converted to NIfTI images using the dcm2niix DICOM to NIfTI converter (The Neuropsychology Lab, University of South Carolina, SC, USA) in this study.

The NIfTI data were preprocessed via the Configurable Pipeline for the Analysis of Connectomes (Craddock et al., 2013) using the default configuration of version 1.7.0. ROI-mean time series data were computed from the Automated Anatomical Labeling (AAL) and the Harvard-Oxford (HO) atlases for the analysis. Moreover, the multi-subject dictionary learning (MSDL) (Varoquaux et al., 2011), Destrieux (Destrieux et al., 2010), and Yeo (Yeo et al., 2011) atlases were added to further examine the inter-atlas differences.

### 2.2. Age prediction modeling

#### 2.2.1. Computational resources

This study was carried out using the TSUBAME3.0 supercomputer at the Tokyo Institute of Technology, Tokyo, Japan. We used this environment interactively via a Python-based web service called Jupyter Lab with a graphics processing unit computing resource for *PyTorch* as a framework for deep learning. All codes used for computing are available at the following URL: https://github.com/yutads/shimane.

#### 2.2.2. Types of age prediction modeling

##### 2.2.2.1. Overview of features

In this study, we carried out multiple types of modeling by aggregating varied features to investigate the best configuration for predicting brain age from resting-state fMRI. These features are classified into the following three levels: *temporal, spatial*, and *architectural*. Here, we interpret the term “feature” in a broader sense, which is not limited to the technical meaning in the field of machine learning.

First, the *temporal* features were implemented as two time series data extraction models, generating a)one matrix for ROI-wise BOLD time series signal change throughout one whole session (run) and b) another for the dynamically changing temporal correlation across the ROI within each time segment (window) stepping forward within the session (run). In this article, a) is abbreviated as the “ROI model” and b) as the “dFC model”.

Next, the *spatial* features were associated with the selection of a brain atlas, which was crucial for parceling the cortices into separate regions where the voxel-wise data were to be averaged. The atlases were classified into anatomical and functional types; the former is principally based on the affinity which is observed at the level of cortical morphology on structural MRI, whereas the latter consists of grouping the areas that show, regardless of physical distances, functionally synchronized responses in various fMRI experiments. We mainly used the former–the AAL or HO–for exhaustive modeling with different feature patterns, but tried to compare these two different types within a limited number of controlled feature conditions.

Last, the *architectural* features involved in our study were in the form of three RNN models utilized in deep learning: the Transformer, Multi-task Transformer, and unilateral or bilateral LSTM. These modeling algorithms are considered suitable for any serial data expressing time-wise relations. Further details are described below in the section on *architectural* features.

##### 2.2.2.2. Temporal features: time series data extraction models

The temporal features used in this study were obtained by the following processes. We obtained the average time series of BOLD signal change for each ROI in each session (run) as the first data extraction method. We regard this method with its outcomes as one of the features concerning data extraction, and refer to it as the ROI-wise mean BOLD lump-sum time series model, or simply as the “ROI Model”. In this setting, we did not compute the time course BOLD correlation between the ROIs while inputting the specifications of our RNN models. For the ROI Model, we used the ROI-wise BOLD signal changes just as they were, solely configuring for inputting the target scope with a whole session (run) of several minutes taken as a single chunk (Figure 1).

**Figure 1.**
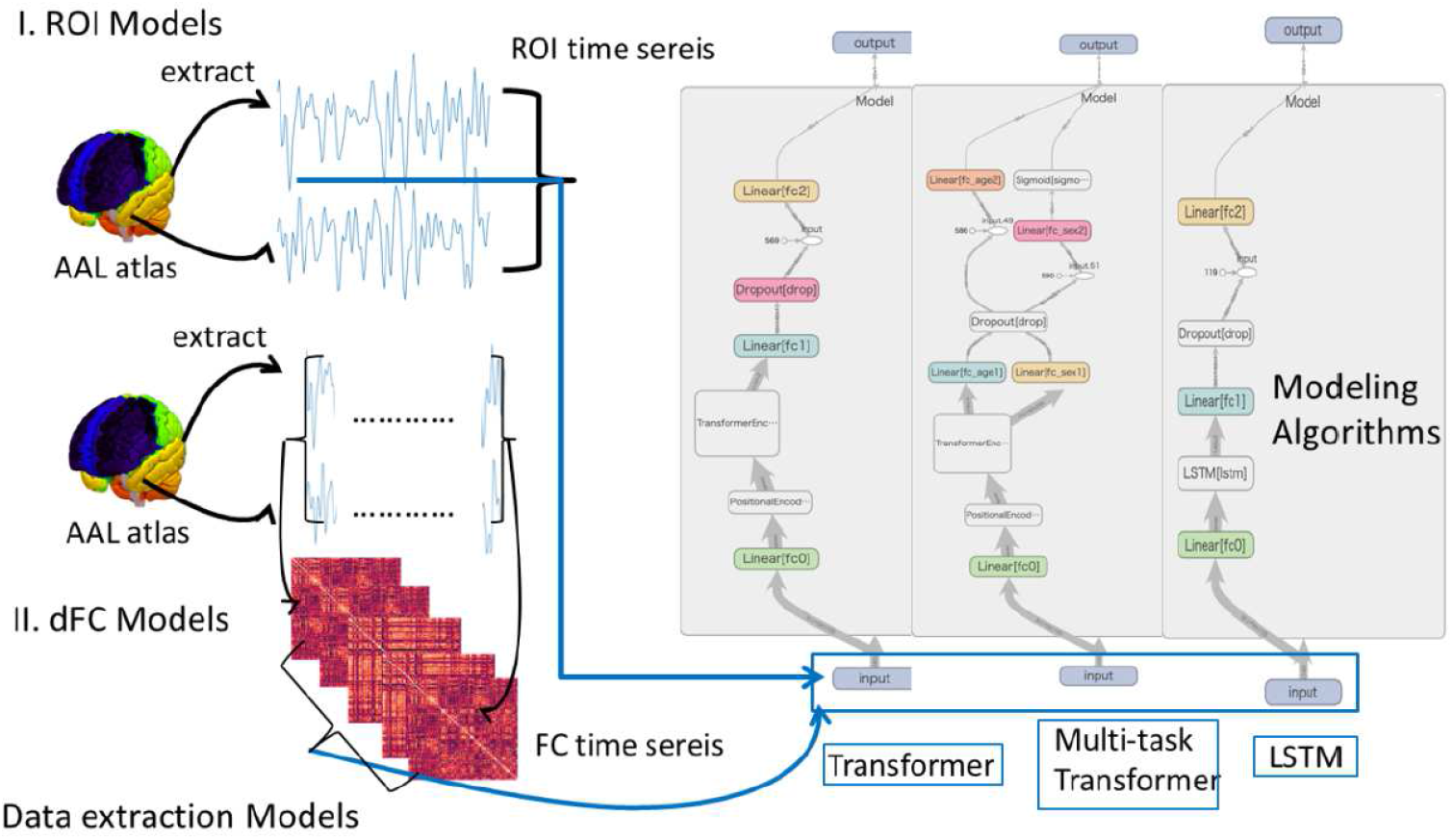
Data extraction models for generating the inputs for the three deep learning algorithms used in this study. The left part represents the two data extraction models, that is, the ROI Model based on the lump-sum ROI-wise mean BOLD time series data (above) and the dfC Model based on the functional connectivity time series using a sliding time window (below). The right part of this figure, sketched using TensorBoard (TensorFlow’s visualization toolkit), shows the three modeling algorithms: Transformer (left), multi-task Transformer (middle), and LSTM (right) with the computational flow architectures.

Next, apart from the ROI Model, we created from the whole session (run) data the so-called dynamic rsFC (abbreviated hereafter as dFC) as a second feature. The data inputs for the dFC were obtained by sliding step by step a window (scope of time zone) over the whole session (run) to compute the ROI-wise temporal correlation within each window position (state). In this sense, the dFC Model can be considered as the data extraction method using a functional connectivity (FC) time series (Figure 1). We assigned a window length of 45 seconds, and the overlapping ratio between the proximate window states to 0.94, which could be considered sufficient time to capture the nonstationarity of FC (Gonzalez-Castillo et al., 2015; Zalesky & Breakspea, 2015).

These two features for data extraction were designed to systematically compare model performances, which was why the ROI Model did not expressly take any correlation matrix built on the ROI-wise BOLD signal change. The ROI Model and dFC Model were comparable based on the similarity in the time-sequence setting. The former took the raw data matrices of ROI by time points in which the values related to activation were recorded in order, while the latter was represented by the matrices of ROI by correlation coefficient values in due course for each window state.

To examine the robustness of the RNN-based models with different parameter values, four age prediction models were created, two for each of the two features of the ROI and dFC Models. For the Transformer-based ROI and dFC data analysis, we created two models (Models 1 and 2) with partially different parameter values. These values were arbitrarily chosen for the value of an example, such that the dimensionality of the input vector (*d_model*) was set to 256 and 128 and the size of the hidden state vector (*hidden_dim*) was set to 1024 and 128, for ROI Models 1 and 2, respectively. We attempted to run the dFC Models 1 and 2 (hereafter abbreviated as dFC 1 and dFC 2, respectively) with two values (0.3 and 0.5, respectively) for attention drop rate (*pos_drop*). For the LSTM-based ROI and dFC Models, two types of variations were prepared (unidirectional and bidirectional). These variations used only the atlases of AAL and HO for the present study, except to make comparisons between atlases when switching the choices of the hyperparameter values and the features.

The dimensions of each of the ROI and dFC Models were as follows. For the AAL atlas, the ROI Model started from a 140 (time points) × 116 (AAL ROIs) matrix for each participant, whereas the dFC computation presupposed a dimensionality of 123 (window states) × 6670 (combination of AAL ROIs), since the ROI-wise correlation coefficient was calculated according to window size. Similarly, the ROI Model data matrix for the HO atlas had dimensions of 140 (time points) × 110 (HO ROIs), while the dimensionality of the dFC matrix was 123 (window states) × 5995 (combination of HO ROIs). These data matrices were used to create models based on the Transformer, LSTM, and multi-task Transformer algorithms.

##### 2.2.2.3. Spatial features: brain atlases

In assessing spatial features, the AAL and HO atlases were chiefly used as structural atlases when combining the other feature patterns in our modeling analysis. These atlases were prepared beforehand in the C-PAC default pipeline setting so that we only had to activate the option of time series extraction in the pipeline .yml file. For inter-atlas comparisons, three other atlases (MSDL (Varoquaux et al., 2011), Destrieux (Destrieux et al., 2010), and Yeo (Yeo et al., 2011)) were adopted in addition to the AAL and HO atlases. The MSDL atlas was used to assess the rsFC dataset of 20 subjects. The Destrieux atlas is characterized by an exact parcellation of cortical surfaces after the meticulous unfolding of each sulcus. The Yeo atlas was created from the rsFC dataset of 1000 subjects, and comprised 7 or 17 regions representing the intrinsic FC networks. In this sense, the Yeo atlas was the most typical functional atlas. These atlases were taken from the module of “datasets” for atlas storage in Nilearn (https://nilearn.github.io/), a well-known Python package for executing machine learning of neuroimaging data. The functions of NiftiMapsMasker (MSDL) or NiftiLabelsMasker (Destrieux and Yeo) prepared in the module of “input_data” were applied to the functional images standardized in the default pipeline of C-PAC using Advanced Normalization Tools.

#### 2.2.3. Architectural features: modeling algorithms

##### 2.2.3.1. Transformer modeling

Transformer is a deep learning algorithm that can handle time series data. This modeling algorithm employs an attention mechanism, made of the queries Q, the keys K, and the values V as main components, and has won many state-of-the-art awards in the field of natural language processing research (Vaswani et al, 2017). Transformer can model long-term dependencies better than LSTM thanks to the attention mechanism (Bahdanau et al., 2014). This RNN algorithm used here, in addition to the Linear Layer, Positional Encoding, and Transformer Encoder Layer, further includes the Dropout Layer and the ReLU function as the activation function. This computational architecture makes it possible to learn the long-range dependencies of series data (Figure 1).

The network structure common to the four age prediction models using Transformer is a feature transformation of the input data, as a third-order tensor, through a Linear Layer for each time axis. This transformation produces a new third-order tensor of the time series. In the next layer for Positional Encoding, time series information is added to the data that has passed through the Linear Layer. The next two Transformer Encoder Layers extract information from the features to which time series information has been added by Positional Encoding. Finally, age prediction is performed by passing the data through two Linear Layers. The mean squared error (MSE) is used to arrive at the loss function in Transformer computing. In this study, the loss was defined as the average of the individual square loss MSE_*n*_ between the predicted age of a person and his/her actual age.

##### 2.2.3.2 Multi-task Transformer modeling

Few studies exist on fMRI analysis using multi-task learning. Huang et al. (2020) proposed a multi-task learning model for auxiliary diagnosis of autism spectrum disease and attention deficit hyperactivity disorder using fMRI images, which achieved reliable classification performance.

Our Multi-task Transformer model adds a partial network to the ordinary Transformer described above to solve the gender classification problem as an auxiliary task in the final layer. We used a Linear Layer to branch the network, which enabled us to solve the gender classification problem and predict age from the Transformer Encoder Layer. Moreover, a sigmoid function was added to the output layer of the gender classification network as an activation function (Figure 1).

The Multi-task Transformer model used for the loss function the weighted sum of both the MSE for age prediction mentioned above and the binary cross-entropy (BCE) to learn binary gender classification. The BCE here is the average across all the BCE losses computed for a participant’s predicted gender and his/her real gender. The losses during the training are shown below (1), where ***x*** represents observed labels and ***y*** represents predicted outcomes. Throughout the cross-validation (CV) which we will discuss hereafter, the loss functions were the same during verification and the training steps.

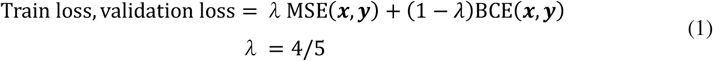

In this study, label smoothing (Szegedyet al., 2019) was introduced in the loss function for the purpose of regularization, since simple multi-task learning tends to cause overlearning. Label smoothing for the learning at the individual level involved a process *n* in which we put *y*_*n*_ = *γ* when the correct label was *y*_*n*_ = 0 and *y*_*n*_ = 1 − *γ* when *y*_*n*_ = 1 held true. In this study, we set *γ* = 0.3 for training only and did not introduce label smoothing at the step of validation in the nested CV which will be discussed below.

##### 2.2.3.3. LSTM modeling

The LSTM model is a deep learning model that can handle time series data, and was first introduced by Hochreiter and Schmidhuber (1997). In our LSTM model, a new third-order tensor of the time series was obtained, followed by the next LSTM layer that extracted the time series information, and finally, age was predicted by passing through two Linear Layers. We adopted both unidirectional and bidirectional LSTM models for each ROI Model and dFC Model as features of time series data extraction.

The unidirectional LSTM learns hidden vectors by following an updated formula, while the bidirectional LSTM performs the updating process for the reverse direction, and merges them with the hidden vectors. This allows for the incorporation of information in both directions. Since this study used a multilayer LSTM model, the input 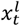 at the *l*^th^ layer becomes a hidden vector 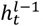, on which dropout is performed. Similar to the Transformer model, only the MSE was used for losses in the LSTM model for age prediction.

#### 2.2.4. Details of learning

For all the models applied in this study, AdamW (Loshchilov & Hutter, 2019) was used for mini-batch learning. AdamW is acknowledged as an improved version of the optimization algorithm called Adam (Kingma & Ba, 2017). We set the batch size to 32, which is the number of data sampled in mini-batch learning. One thousand epochs were arranged for learning on ROI Model data, and 700 epochs for learning on the dFC Model. The same model was used for both the AAL and HO atlases in the analysis. In this study, the default PyTorch settings were employed as the hyperparameters of AdamW, except the initial learning rate which was set to 1e − 4. Furthermore, early stopping was introduced into this analysis. Training was terminated when the loss value for validation could not be reduced further after 200 consecutive epochs. The model that was able to minimize the loss of data for validation in all cases was adopted in this study.

#### 2.2.5. Evaluation methods

By way of accuracy assessment, the following CV and prediction accuracy calculation methods were used. In this study, CV was performed via nested CV. First, the overall data were divided into 5 parts, and 4/5 of the divided data was used as Training data and 1/5 as Test data. The specific method of division used during iteration is shown in Figure 2. The loop used in the process of first dividing the data into Training and Test data was called the outer loop, and the loop for the process of further dividing the Training data into (sub) Training and Validation data was called the inner loop.

**Figure 2.**
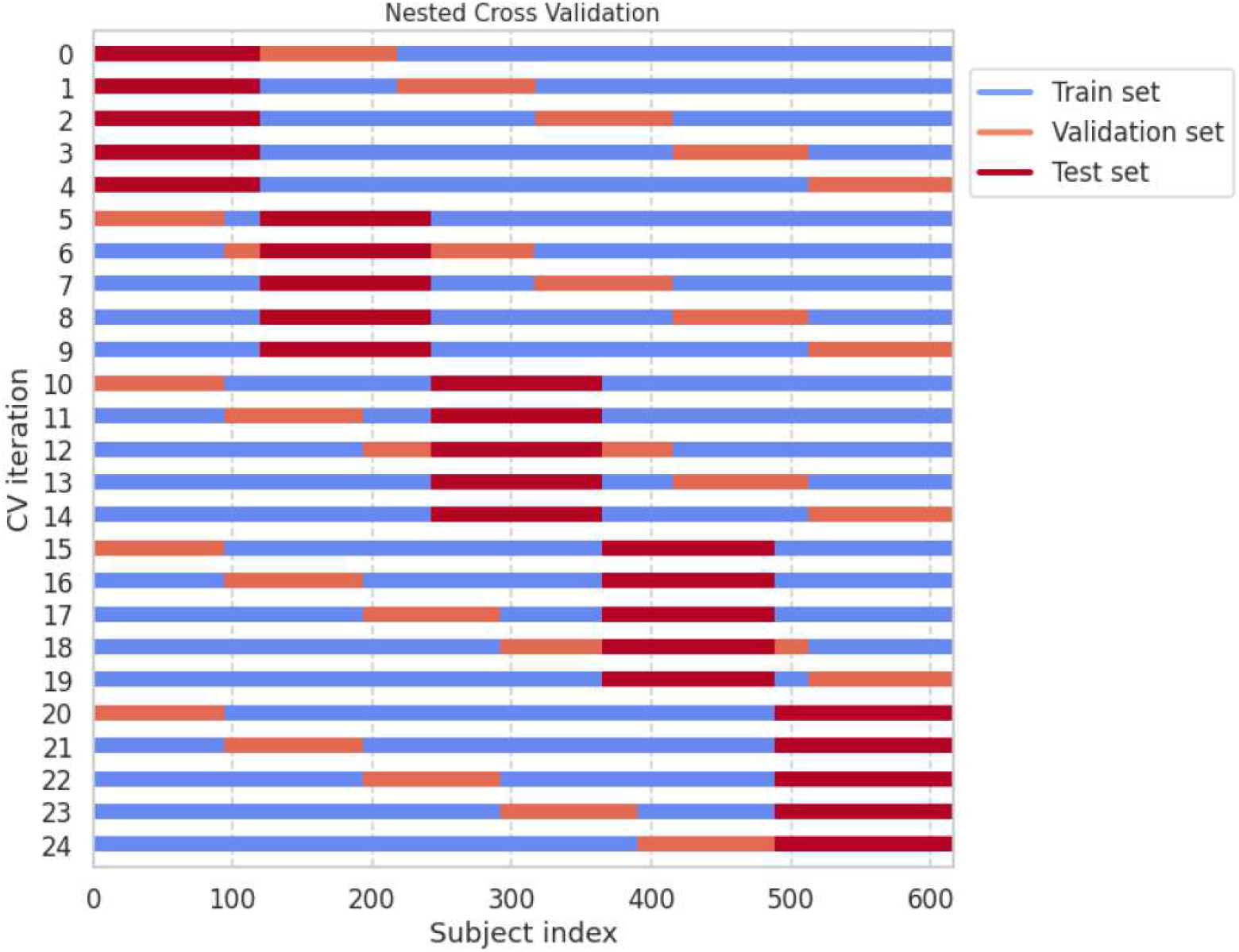
Outer and inner loops of the nested CV configured for evaluating our models. The horizontal axis represents the subject index, whereas the vertical axis represents the process step number in the CV iteration. Each color of the three data subsets (training set, validation set, or test set) represents a subject assessed in each step of the nested CV.

To evaluate the results of CV, the mean absolute error (MAE) was used as a measure of accuracy in age prediction (2). Here *x*_*n*_ represents a participant’s predicted age and *y*_*n*_ his/her actual age. The calculated MAE_*n*_ values were averaged for all participants to output the MAE(*x,y*) as the predictive index for the results of this study.

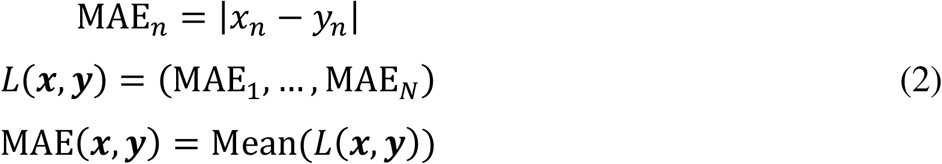

#### 2.2.6. Bias adjustment

With regard to age prediction using deep learning, it has been widely acknowledged that the raw output is sometimes characterized by overestimation for younger age groups and underestimation for older age groups. Consequently, the regression coefficient of the relationship between real and predicted ages is substantially lower than 1, and the regression lines considerably tilt down. A previous study showed that this overestimation or underestimation is not unique to a particular age group, and that regardless of data or methods, it is hard to remove this form of bias by including a uniform age distribution (Liang et al, 2019). In this study, therefore, bias correction was used in some cases. Two techniques of bias correction were introduced by de Lange and Cole (2020). The first technique recycles the Validation set to store the slope and intercept to be applied to the Test set as expressed in equation (3). Since real age is not used in this formula, bias correction can be performed without causing information leakage.

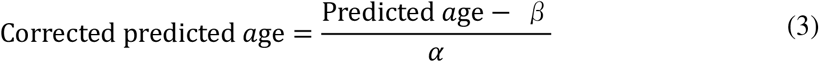

Note that the second technique uses actual age for re-computation, so it is intended only to correct for bias, not to predict age in the original sense. It consists of two steps that adopt the least-square method to the Test set to regress the predicted age on the actual age, and corrects the raw predicted age using the slope and the intercept of the regression line as shown in equation (3).

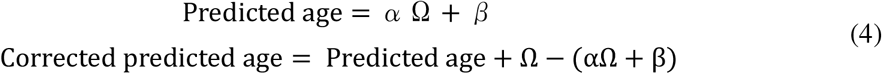

In this study, the first bias correction technique overestimates the MAE relative to the second since it does not use actual age, but it is considered faithful to the original prediction, and effective because of the invariability of the correlation coefficients. In particular, the second technique is able to account for the potential age dependence of the target variables. However, when evaluating the accuracy of each technique, comparisons should be made with the results obtained before any correction was made. See the Supplementary Materials, Addendum 1 for details on the results of bias correction.

## 3. Results

### 3.1. Output of each model

In this study, three *architectural* models based on the Transformer, Multi-task Transformer, and LSTM algorithms were created for comparison, and age was predicted using each of these deep learning models. Figure 3 illustrates the relationship between predicted and actual ages determined using the Transformer or LSTM models with the AAL or HO atlases (*spatial* features). In this figure, eight modeling conditions were attempted for each atlas or modeling algorithm (Transformer or LSTM). Thus, a set of four panels was created for the combination of atlases and modeling algorithms, and within each panel, either the ROI Model or dFC Model (*temporal* features) was tested by changing hyperparameter values (Transformer) or network directionality (LSTM) (numbered Model 1 and Model 2).

**Figure 3.**
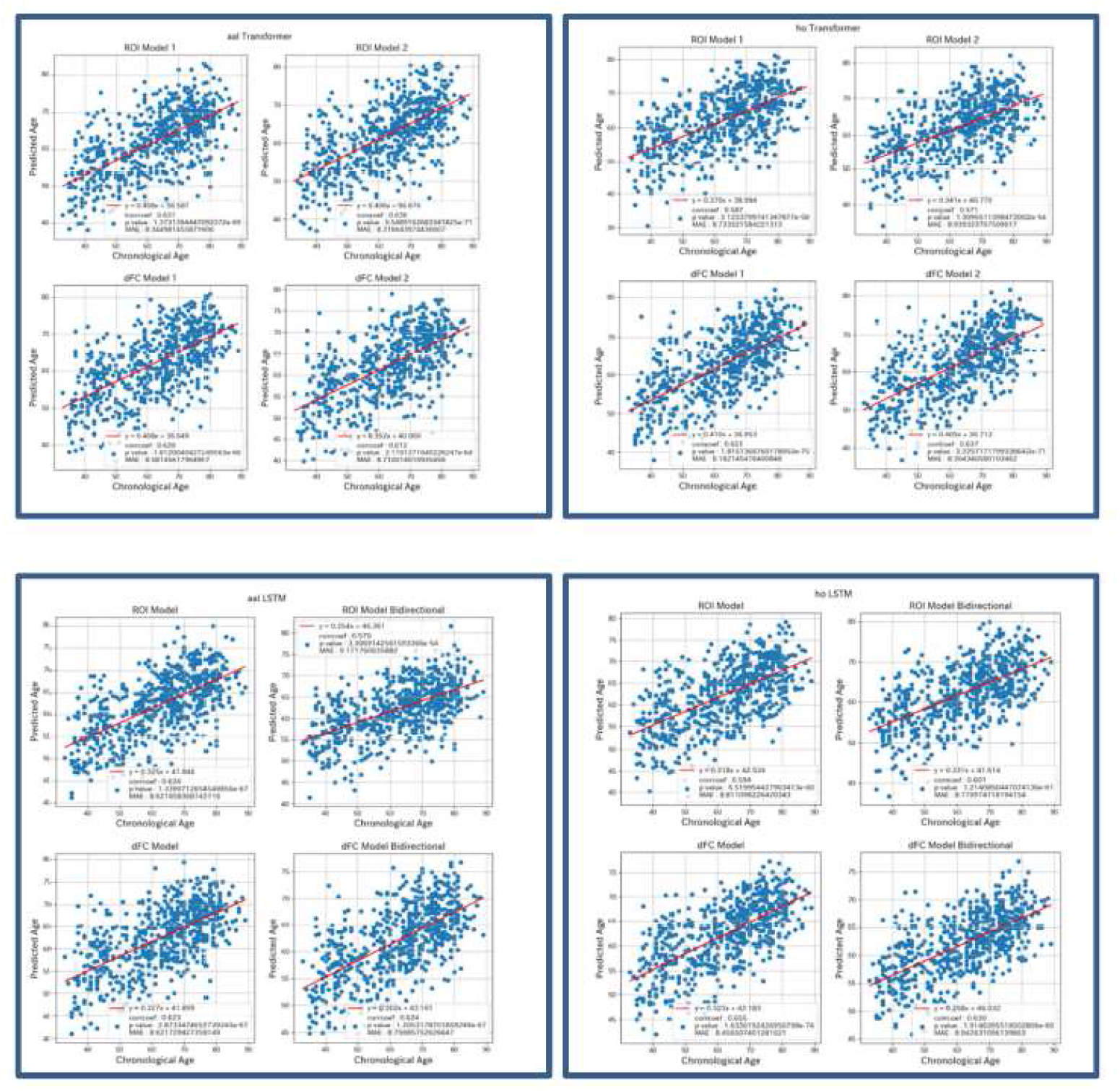
Relationships between predicted and actual ages determined using the Transformer or LSTM algorithms with the AAL or HO atlases (spatial features). Above left: Relationship between predicted and actual ages determined using the Transformer model with the AAL atlas. Above right: Relationship between predicted and actual ages determined using the Transformer algorithm with the HO atlas. Below left: Relationship between predicted and actual ages determined using the LSTM algorithm with the AAL atlas. Below right: Relationship between predicted and actual ages determined using the LSTM algorithm with the HO atlas. In each graph, the horizontal axis represents chronological (real) ages and the vertical axis represents predicted ages. Each dot represents an individual participant.

For each of the results, the P-values of the correlation coefficients for the AAL atlas were on the order of the minus 60th to the 70th power of 10, indicating that predictions were successfully made for this setting. Likewise, for the HO atlas, the P-values of the correlation coefficients were on a similar order, indicating similarly good performance. Table 1 represents the MAEs for the differences between predicted and actual ages determined using the Transformer or LSTM algorithm. Table 2 shows the slopes of the linear regression lines drawn for predicted age as a dependent variable and actual age as an independent variable when using the Transformer or LSTM algorithms. The slopes which were substantially below 1.0 in this study were due to the biased sample from middle-aged and elderly groups, which was inevitable given the practical constraints of the Brain Dock. Those values are indeed far from being on a par with those of prior studies in which age group distribution could be perfectly manipulated at the experimental design stage, but the MAE values were not that much larger compared to those of similar studies.

**Table 1.** MAEs for the differences between predicted and actual ages determined by the Transformer or LSTM algorithm.

| Normal | ROI 1 | ROI2 | dFC 1 | dFC 2 |
| --- | --- | --- | --- | --- |
| Transformer AAL | 8.345 | 8.317 | 8.581 | 8.71 |
| Transformer HO | 8.733 | 8.939 | 8.182 | 8.304 |
| LSTM AAL | 8.622 | 9.172 | 8.622 | 8.757 |
| LSTM HO | 8.811 | 8.771 | 8.457 | 8.943 |

MAE: mean absolute error; ROI 1, ROI 2: ROI average time series Models 1 and 2; dFC 1, dFC 2: dynamic functional connectivity Models 1 and 2. The hyperparameter value settings were different between Model 1 and Model 2.

**Table 1.** Slopes of the linear regression lines of the relationship between predicted and actual ages determined by the Transformer and LSTM algorithms.

| Normal | ROI 1 | ROI 2 | dFC 1 | dFC 2 |
| --- | --- | --- | --- | --- |
| Transformer AAL | 0.408 | 0.406 | 0.408 | 0.352 |
| Transformer HO | 0.37 | 0.341 | 0.41 | 0.405 |
| LSTM AAL | 0.325 | 0.254 | 0.327 | 0.302 |
| LSTM HO | 0.318 | 0.331 | 0.323 | 0.258 |

MAE: mean absolute error; ROI 1, ROI 2: ROI average time series Models 1 and 2; dFC 1, dFC 2: dynamic functional connectivity Models 1 and 2. The hyperparameter value settings were different between Model 1 and Model 2.

From the viewpoints of learning architectures, the results showed that the differences between the modeling algorithms were not so much more pronounced than expected. The Transformer model was slightly more accurate in terms of the MAE than the LSTM; however, the total difference in MAE across the eight conditions noted in Figure 3 was not significant (mean for Transformer: 8.514, mean for LSTM: 8.770, P = 0.055). However, the slopes of the resulting regression lines were constantly higher for the former than the latter (mean for Transformer: 0.388, mean for LSTM: 0.305, P = 7.81e-05). Thus, we could hypothesize that the Transformer algorithm has a slight advantage over the LSTM algorithm in this context.

On the other hand, the results based on the Multi-task Transformer algorithm were worth commenting upon. Figure 4 shows the relationship between predicted and actual ages determined using this modeling algorithm. The panels were distinguished according to the choice of atlas (*spatial* features), and each of them included four modeling results based on either the ROI Model or the dFC Model (*temporal* features), and either one hyperparameter value setting or another (Figure 3). Interestingly, the combination of the AAL and ROI Model plus that of the HO and dFC Model always recorded better accuracy in terms of MAE than the opposite cross combination pattern. However, any apparent assumption concerning this may lead to incorrect conclusions except the existence of important variance across the atlas-based ROI settings. In comparing the *architectural* features between the performances of the Multi-task Transformer and the original Transformer algorithms (Tables 1, 2, 3, and 4), there was no significant difference in prediction accuracy (mean MAE for Transformer: 8.514, mean MAE for Multi-task Transformer: 8.627, P = 0.377; mean slope for Transformer: 0.388, mean slope for Multi-task Transformer: 0.358, P = 0.053).

**Figure 4.**
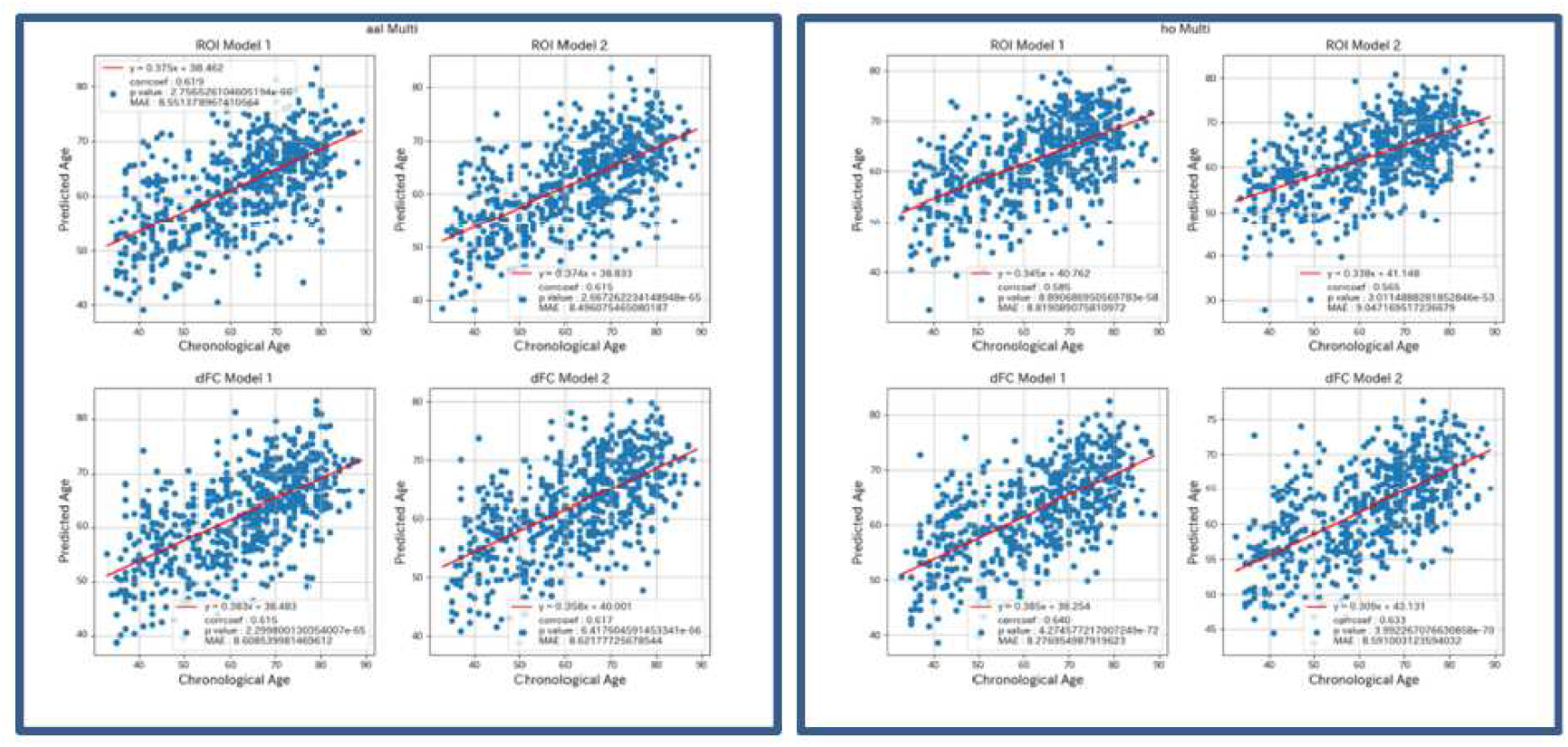
Relationships between predicted and actual ages determined using the Multi-task Transformer algorithm for ROI Models 1 and 2 and dFC Models 1 and 2 with the different atlases. Left: AAL; Right: HO. In each graph, the horizontal axis represents chronological (real) ages, whereas the vertical axis represents predicted ages. Each dot represents an individual participant.

**Table 3.** MAEs for the differences between predicted and actual ages determined via multi-task learning.

| Multi_task_Normal | ROI 1 | ROI 2 | dFC 1 | dFC 2 |
| --- | --- | --- | --- | --- |
| Transformer AAL | 8.551 | 8.496 | 8.609 | 8.622 |
| Transformer HO | 8.819 | 9.047 | 8.277 | 8.591 |

MAE: mean absolute error; ROI 1, ROI 2: ROI average time series Models 1 and 2; dFC 1, dFC 2: dynamic functional connectivity Models 1 and 2. The hyperparameter value settings were different between Model 1 and Model 2.

**Table 4.** Slopes of the linear regression lines of the relationship between predicted and actual ages determined via multi-task learning.

| Multi_task_Normal | ROI 1 | ROI 2 | dFC 1 | dFC 2 |
| --- | --- | --- | --- | --- |
| Transformer AAL | 0.375 | 0.374 | 0.383 | 0.358 |
| Transformer HO | 0.345 | 0.338 | 0.385 | 0.309 |

MAE: mean absolute error; ROI 1, ROI 2: ROI average time series Models 1 and 2; dFC 1, dFC 2: dynamic functional connectivity Models 1 and 2. The hyperparameter value settings were different between Model 1 and Model 2.

Furthermore, the aforementioned bias corrections were computed, generating regression lines with slopes whose values were almost 1 for the relationship between real and adjusted predicted ages. The MAEs for evaluating the models were corrected by using the adjustment techniques 1 and 2 recorded an average of 14.44 for the former and 4.94 for the latter, with a mean difference of 9.51 (P = 2.28e-11), across the 8 combinations of features and hyperparameters. The details are discussed in the Supplementary Materials (Figure A1, Tables A1–A4). Regarding the results of gender discrimination in this multi-task learning, we obtained a solution within the RNNs whose main purpose was extraneous to the binary classification but related to outputting the real number values for age prediction. For the sake of comparison, we created another model that only performed gender discrimination without age prediction (defined as single-task prediction); however, this did not show much difference in accuracy from the multi-task learning results. The accuracy rates averaged across the four conditions (2 hyperparameter settings by 2 data extraction models by 2 atlases) were as follows: multi-task model: 0.58, single-task model: 0.65. The details of this model are also presented in the Supplementary Materials (Figure A2, Table A5).

### 3.2. Comparison between modeling conditions

In this study, we trained multiple deep learning models for age prediction or gender discrimination with the ROI and dFC Models as *temporal* features selecting atlases as *spatial* features for determining brain parcellation. To know in detail the relativeness of these models, we produced scatter plots to compare the predicted results for each subject across hyperparameter value settings, various feature conditions, and modeling algorithms (*architectural* features). The magnitudes and P-values of the correlation coefficients in these plots were also calculated to evaluate the affinity between predicted results. Consequently, we found that the differences in correlation coefficients were largely due to *spatial* features (or *temporal* features to a lesser extent) rather than due to modeling algorithms.

The two scatter plots in the top left panel of Figure 5 were arranged to compare the predicted results of different modeling algorithms (Transformer, LSTM, and bidirectional LSTM) for the same features. The correlation coefficient was inferior but very close to 0.9. For the two plots in the upper right panel, the left one showed the correspondence between the predicted results of the two Transformer models with different hyperparameter values when the rsFC ROI average time series under the AAL atlas was adopted as a feature condition. The right figure in the same panel was almost identical for the modeling algorithm and atlas, but used dFC with the two hyperparameter value settings. Both cases yielded correlation coefficients larger than 0.95.

**Figure 5.**
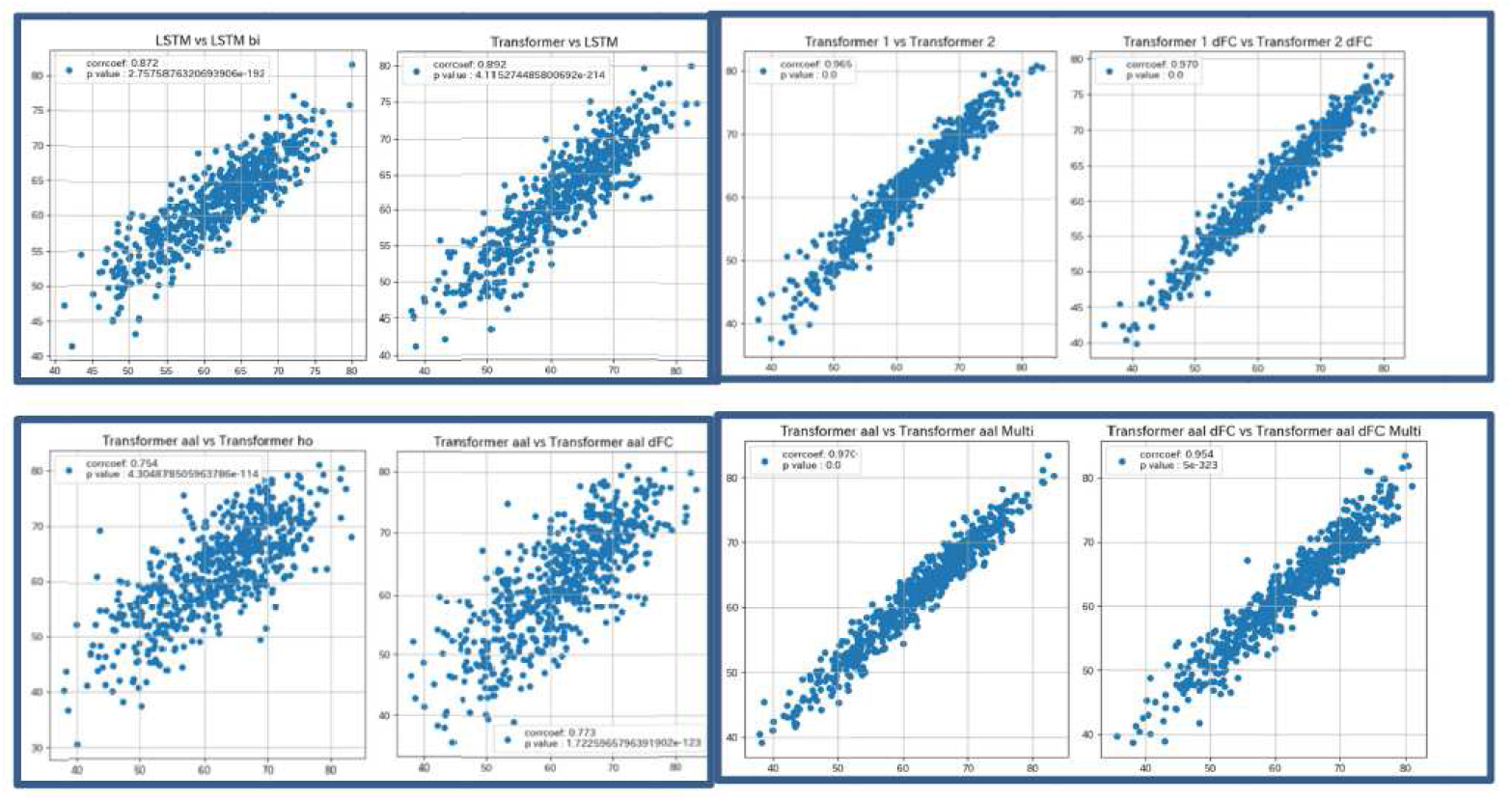
Comparison of predicted results obtained by changing hyperparameter settings, data extraction methods, deep learning algorithms, and brain atlases. Above left: Comparison of predicted results between different modeling algorithms under the same feature conditions (LSTM vs bidirectional LSTM, and Transformer vs LSTM). Above right: Comparison of predicted results between the same models with different hyperparameter values but under the same feature conditions (ROI Model 1 vs ROI Model 2, and dFC Model 1 vs dFC Model 2). Below left: Comparison of predicted results between the same model (algorithm) but under different feature (atlas or data extraction model) conditions (AAL vs HO, and ROI Model vs dFC Model). Below right: Comparison of predicted results between the modeling algorithms with and without multi-task learning under the same (atlas or data extraction model) features. In each graph, the horizontal axis represents chronological (real) ages, whereas the vertical axis represents predicted ages. Each dot represents an individual participant.

The panel including the two plots in the lower left shows a comparison between the predicted results of the same modeling algorithm for different features (*temporal* or *spatial*). The correlation coefficient of the relationship between the AAL and HO atlases as *spatial* features for the Transformer algorithm was approximately 0.754 (left). On the right side is a plot of the results of Transformer prediction with the same atlas (AAL) but with different time series data extraction models of the ROI time series or dFC. The correlation coefficient was 0.773 (right). Finally, the two plots in the lower right panel were created to compare the predicted results for the same features (*temporal* or *spatial*) but with or without multi-task learning, and showed little difference between them (both larger than 0.95).

### 3.3. Comparison between atlases

In this section, we introduce three extra atlases: MSDL (Varoquaux et al., 2011), Destrieux (Destrieux et al., 2010), Yeo-thick-7, and Yeo-thick-17 (Yeo et al., 2011), in addition to the AAL and HO atlases, to predict age and to evaluate these predictions. In applying the Transformer algorithm to ROI and dFC Models, the two hyperparameter settings were employed as shown in Model 1 and Model 2; thus, four results were paneled per atlas (Figure 6). The results of correlation were indeed all significant but with a considerable difference across atlases (Figure 6). For example, the correlation coefficients of the relationships between real and predicted ages were 0.505 (MSDL), 0.605 (Destrieux), 0.295 (Yeo-thick-7), and 0.462 (Yeo-thick-17) for ROI Model 1 determined via the Transformer algorithm. The slopes of the regression lines were 0.267 (MSDL), 0.319 (Destrieux), 0.145 (Yeo-thick-7), and 0.271 (Yeo-thick-17) under the above-mentioned conditions (Table 5). It is worthwhile here to remark on the difference across the atlas types which were anatomical or functional and relationships as well. The Destrieux atlas is a typical product of anatomical parcellation for delineating gyral and sulcal regions of the cortical surfaces; it falls under the same category as the AAL and HO atlases, which yielded comparable outcomes. The MSDL, Yeo-thick-7, and Yeo-thick-17 atlases are based on the functional parcellation composed of FC networks; thus, the numbers of regions are small (38, 7, and 17, respectively). The results for these functional atlases were less accurate than the structural ones due to segmentation granularity (Figure 6). In addition, they might have some issues with robustness in terms of variability in the results of linear regression (Table 5). However, it is noteworthy that all the functional atlases recorded highly significant accuracy with fewer than 50 regions and even less than 10.

**Figure 6.**
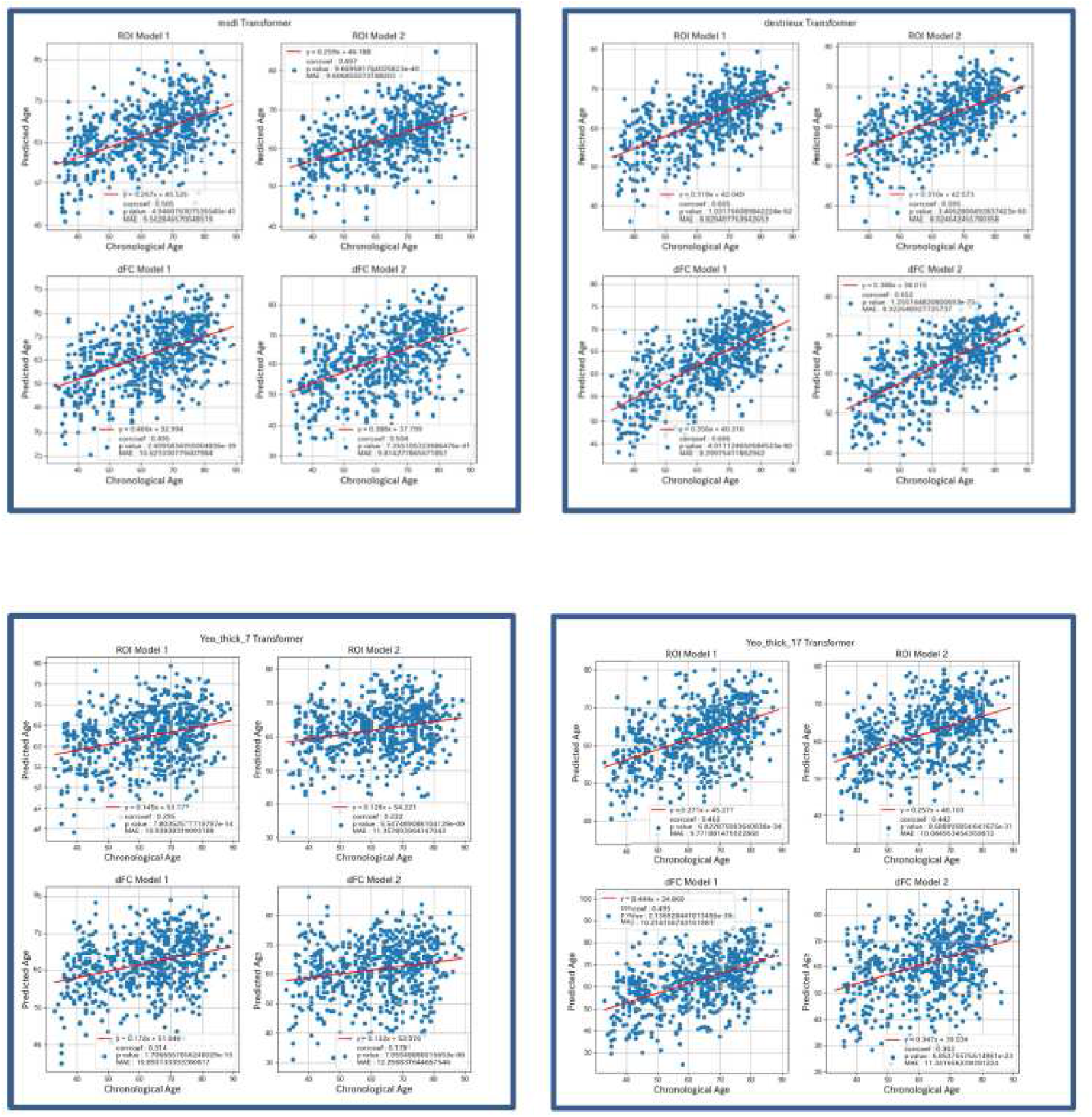
Comparison of predicted results obtained using the different atlases with the Transformer algorithm for the ROI and dFC Models. Above left, MSDL; above right, Destrieux; below left, Yeo-thick-7; below right, Yeo-thick-17. In each graph, the horizontal axis represents chronological (real) ages, whereas the vertical axis represents predicted ages. Each dot represents an individual participant.

**Table 5.** Slopes of the linear regression lines of the relationship between predicted and actual ages determined with the Transformer algorithm according to atlas selection.

| Transformer models | ROI 1 | ROI 2 | dFC 1 | dFC 2 |
| --- | --- | --- | --- | --- |
| MSDL | 0.267 | 0.259 | 0.466 | 0.388 |
| Destrieux | 0.319 | 0.310 | 0.356 | 0.388 |
| Yeo-thick-7 | 0.145 | 0.128 | 0.172 | 0.132 |
| Yeo-thick-17 | 0.271 | 0.257 | 0.444 | 0.347 |
| AAL | 0.408 | 0.406 | 0.408 | 0.352 |
| HO | 0.37 | 0.341 | 0.41 | 0.405 |

ROI 1, ROI 2: ROI average time series Models 1 and 2; dFC 1, dFC 2: dynamic functional connectivity Models 1 and 2. The hyperparameter value settings were different between Model 1 and Model 2.

## 4. Discussion

The correlation coefficient of the relationship between ages predicted by two models, which differed only in terms of hyperparameter settings, was very high (approximately 0.975), under the same feature conditions (identical modeling algorithms, data extraction methods, and atlases for identifying brain areas). This stability argues for the robustness of age prediction modeling with respect to value assignment. Next to that, the correlation coefficient for the two models that differed in terms of modeling algorithms (*architectural* features) under the same *temporal* and *spatial* feature conditions (identical data extraction methods and atlases) was close to 0.9. On the other hand, the correlation coefficient dropped to less than 0.8 when there was a difference in *spatial* or *temporal* feature, that is, when we only changed either the data extraction method or the atlas, applied the same modeling algorithm, and kept the other feature for control. This indicated that the *temporal* or *spatial* feature selected in advance has a larger effect on the model than hyperparameter settings and modeling algorithms. Based on the above, we can safely assert that viewed from the comparative stability in correlation across models, there was relatively little need to refine the modeling algorithms themselves, regardless of differences in other features if they were kept constant.

In this article, we compared the ROI time series framework through an undivided session as a whole with the dFC framework in which the short-term temporal correlation was computed for each position of the time window which slid in small steps on the progress of a session. A vast body of literature highlights the characteristics of brain age with respect to each of the two data extraction methods. For the former method with a long history of research, the FC maturity index proposed by Dosenbach et al. (2010) might be useful in this field in that it is not only suited to the classical biological model of growth and maturation based on neurophysiological changes. It provided landmark insights showing that with aging, an increase in connectivity is seen within the intrinsic FC networks, whereas a decrease is observed between them. The nature of secular network modulation might play the role of cue for age prediction modeling; however, the dFC that introduces temporal variability into an rsFC time course session recently turned out to be crucial in providing precious information on age-dependent dynamic functional modulation patterns. Chang and Glover (2010) showed that it was meaningful to incorporate time course variability and dynamics into the progress of synchronization of BOLD signal change across brain regions. For example, it was found, by creating an ALFF-FC Map from the dFC and calculating brain maturity scores, that important age-dependent connections exist mainly between intrinsic functional connectivity networks (Qin et al., 2015). Another study that analyzed dynamic switching between dFC states found, using clustering methods, that age correlated with dwelling time among the dFC states showing weak interactions throughout the brain (Tian et al., 2018).

Such metrological diversity suggests that these two data extraction methods could provide a large amount of complementary and non-overlapping information in BOLD time course synchronization. The ROI-wise BOLD signal change patterns that evolve might carry rich information macroscopically (ROI Model) as well as microscopically (dFC). However, according to the results of our study, particular data extraction methods did not particularly contribute to the outperformance in prediction accuracy for almost all the atlases. Considering data size, the ROI-averaged time series data with a single chunk (ROI Models) are likely to be simpler in clinical practice when focusing on age prediction (ROI Model_AAL: 189 Mb, ROI Model_AAL: 181 Mb, dFC_AAL: 3477 Mb, dFC_HO: 3190 Mb as .csv files). In this sense, the data size might be selectively considered for practical clinical usage; thus, the ROI Model would be preferable as a default in medical applications such as the Brain Dock.

When it comes to the choice of the atlas, the difference between the most conventional ones (AAL and HO) was relatively small. However, problems were identified in the other atlases, such as the lower correlation between real and predicted ages. This flaw was expressly found in the resting-state functional connectivity-based atlases because the restriction in the number of networks implied nothing but poor spatial resolution. The choice of atlas is basically decisive, in the sense that care must be taken when including prior findings of functional anatomy in the interpretation of age prediction. It might be worth removing each ROI from the atlas to evaluate its relative importance from the accuracy drop in the result. In short, atlas selection is of paramount importance in the standardization of brain checkup systems, since we will need to further assess the information from anatomical scans. However, it cannot be reckoned that the functional atlases, namely the MSDL and Yeo atlases, recorded accuracy on a level with the anatomical ones when given odds for a modest number of regions. This finding corroborates the statement made by Qin et al. (2015) that senile changes in the brain are observed primarily at the level of intrinsic FC networks.

In this paper, we attempted to integrate gender guessing into age prediction modeling intermingling regression and discrimination within a single network. The multi-task learning model could provide significant classification accuracy for gender discrimination, although it did not improve accuracy compared to the independent model without multi-task learning. Two possible reasons could be raised. First, contrary to the intuitive premise that gender difference would be relevant to brain senescence, it may not be the case that age and sex information could provide unified complementary modeling. The second reason may be that our model and data conditions were ill-suited to that purpose: they simply did not reflect gender information sufficiently. In this study, the accuracy of the gender discrimination model was at most 70%, which was not sufficiently high compared to previous studies. From the viewpoint of data gathering, restrictions on age variance and availability of health facilities are difficult to lift for more refined experiments if we stay in the context of the conventional Brain Dock. The best scores have been recorded from well-designed fMRI experiments, which demonstrated, for example, 87% accuracy (Zhang et al., 2018) with a large number of subjects (n=820) and narrow age distribution (range, 22–37 years), or 71% accuracy (Satterthwaite et al., 2014) with 674 individuals between the ages of 9 and 22 years. However, since little is known about gender discrimination in fMRI, we cannot rule out the possibility that with a new type of modeling, adding gender information to the model could increase the accuracy of age prediction.

It is worth noting that for the raw models without bias correction, as we described above, the slopes of the regression lines for age prediction came out noticeably lower. This tendency has been known to be universally inevitable, and in the present study, there was no exception in the results of various types of modeling. Bias correction was successfully executed for this Brain Dock dataset, and future research should be fueled to take advantage of correction with the view of drawing clinical inference on an individual’s cognitive reserves from the difference between real age and bias-corrected predicted age (Supplementary Materials, Addendum 1).

The present study had some limitations. Recently, increasing times for image scanning and number of participants have opened the door to growing datasets expected to improve modeling accuracy in machine learning. We should thus envision ideal collections with large amounts of MRI scans for the Brain Dock by overcoming the unavoidable constraints specific to this medical service. Although we were able to obtain high accuracy from the data of 600 individuals acquired at one hospital, this was far from sufficient since most of them were of advanced age. We must address an underlying cause of the flaws in data gathering bias. The Brain Dock service in Japan responds in substance to the demands of health-conscious elderly people, and more or less to those of private companies urging their middle-aged employees to undergo medical check-ups. Consequently, a major challenge for this service is recruiting younger patients and convincing them of the need to begin participating in dementia prevention much earlier than expected. The age distribution ideal for predicting brain age by machine learning will ensue from this effort in improving public awareness for the realization of lifelong health. Combined with this optimization for data collection, another issue should be addressed, which involves investigating a multi-site data harmonization methodology for integrating Brain Dock data from multiple organizations.

## 5. Conclusions

To the best of our knowledge, this study is the first to apply resting-state functional time series modeling to the Brain Dock, a medical system unique to Japan. We aimed, for the first time, to systematically compare the results of age prediction acquired from various deep learning models such as the Transformer, Multi-task Transformer, and LSTM models. A highly significant correlation was observed between actual and predicted ages from all types of methodologies. Robust and stable results recording correlations of more than 0.9 for prediction accuracy were obtained (i) with different hyperparameter value settings, but with the same data extraction methods applied to the same atlas as features as well as the same modeling algorithm (controlling *temporal, spatial*, and *architectural* features). It was also the case when (ii) we used different modeling algorithms (to test the *architectural* features) with the same hyperparameter setting and the same data extraction methods as features based on the same atlas (maintaining identical *temporal* and *spatial* feature conditions). In summary, robust and highly significant levels of correlation were obtained between actual and predicted ages from all types of methodologies. Furthermore, we determined that some conditions have a relatively large impact on prediction performance based on extended comparisons. The correlation for prediction accuracy dropped to 0.75–0.8, when (iii) the same atlas was used (maintaining *spatial* features) but attempted to compare the ROI Model vs the dFC Model as contrasting data extraction models (*temporal* features), or particularly when (iv) different atlases (to test *spatial* features) were used even with the same modeling conditions for *temporal* and *architectural* features. In particular, we found that age prediction was greatly affected by the choice of atlas constructed by nodes of intrinsic functional networks regardless of the number of included areas. Moreover, it turned out that multi-task learning with other phenotype data (related to gender difference) was possible, but did not improve the prediction accuracy as much as expected. Despite these limitations, our results could provide a hopeful prospect of introducing fMRI into the field of neuro-clinical practice.

## Data availability

Limited phenotype data (subject ID, age, and sex) and the inputs (ROI mean time series data based on the AAL and HO atlases only) of the neural networks generated for this study are available with all the Python codes for data analysis at https://github.com/yutads/shimane. However, the functional MRI images are not open to the public through the internet due to the conditions for approval of the study set by the institutional review boards.

## Author contributions

YM: conceptualization, data curation, analysis, investigation, methodology, writing – original draft, writing – review & editing. KO: conceptualization, data acquisition, investigation, methodology, writing – review & editing, supervision. MT: conceptualization, writing – review & editing, supervision. SY: conceptualization, data acquisition, writing – review & editing, supervision. NO: data curation, analysis. HA: conceptualization, data curation, analysis, investigation, methodology, writing – review & editing, supervision. All the authors contributed to the manuscript revision, read, and approved the submitted version. All authors have seen and approved the manuscript.

## Acknowledgements

The authors would like to thank Editage (www.editage.jp) for English language editing.

## Supplementary materials

## Appendix 1: Bias correction

**Figure A1.**
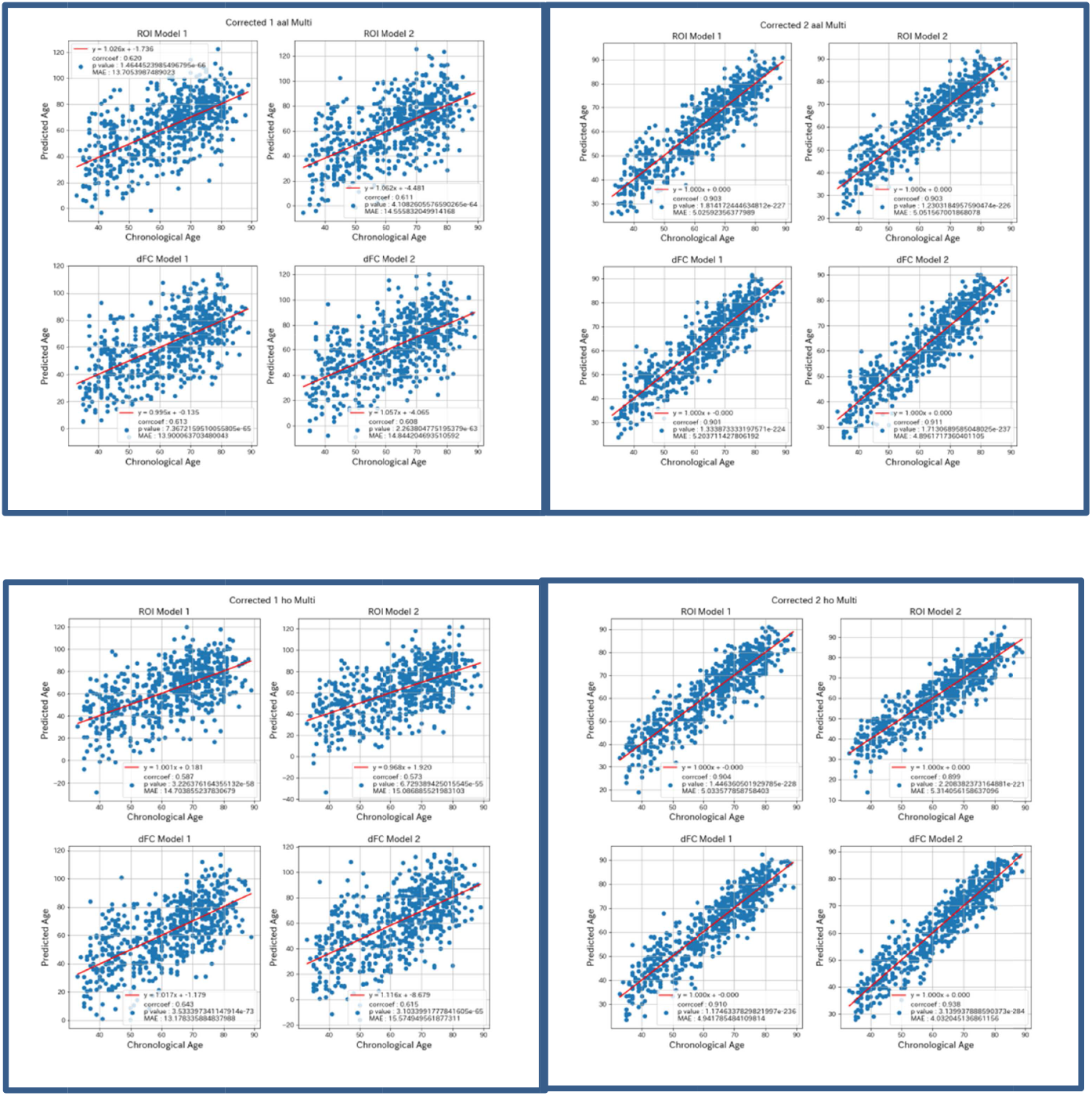
Above left: Relationship adjusted by the bias correction technique 1 between predicted and actual ages using the multi-task Transformer model with the AAL atlas. Above right: Relationship adjusted by the b as correction technique 2 between predicted and actual ages using the multi-task Transformer model with the AAL atlas. Below left: Relationship adjusted by the bias correction technique 1 between predicted and actual ages using the multi-task Transformer model with the HO atlas. Below right: Relationship adjusted by the bias correction technique 2 between predicted and actual ages using the multi-task Transformer model with the HO model. In each panel, the results are demonstrated for ROI Model 1 (above left), ROI Model 2 (above right), dFC Model 1 (below left), and dFC Mode 2 (below right).

**Table A1.** MAEs for the difference adjusted by the bias correction technique 1 between actual and predicted ages adjusted with the multi-task Transformer model

| Multi_task_Corrected_1 | ROI 1 | ROI 2 | dFC 1 | dFC 2 |
| --- | --- | --- | --- | --- |
| Transformer AAL | 13.705 | 14.556 | 13.9 | 14.844 |
| Transformer HO | 14.704 | 15.087 | 13.178 | 15.575 |

MAE: mean absolute error; ROI 1, ROI 2: ROI average time series Models 1 and 2; dFC 1, dFC 2: dynamic functional connectivity Models 1 and 2. The hyperparameter value settings were different between Model 1 and Model 2.

**Table A2.** MAEs for the difference adjusted by the bias correction technique 2 between actual and predicted ages adjusted with the multi-task Transformer model

| Multi_task_Corrected_2 | ROI 1 | ROI 2 | dFC 1 | dFC 2 |
| --- | --- | --- | --- | --- |
| Transformer AAL | 5.026 | 5.052 | 5.204 | 4.896 |
| Transformer HO | 5.034 | 5.314 | 4.942 | 4.032 |

**Table A3.** Slopes of the linear regression lines adjusted by the bias correction technique 1 between predicted and actual ages with the multi-task Transformer model

| Multi_task_Corrected_1 | ROI 1 | ROI 2 | dFC 1 | dFC 2 |
| --- | --- | --- | --- | --- |
| Transformer AAL | 1.026 | 1.062 | 0.995 | 1.057 |
| Transformer HO | 1.001 | 0.968 | 1.017 | 1.116 |

**Table A4.** Slopes of the linear regression lines adjusted by the bias correction technique 2 between predicted and actual ages with the multi-task Transformer model

| Multi_task_Corrected_2 | ROI 1 | ROI 2 | dFC 1 | dFC 2 |
| --- | --- | --- | --- | --- |
| Transformer AAL | 1.0 | 1.0 | 1.0 | 1.0 |
| Transformer HO | 1.0 | 1.0 | 1.0 | 1.0 |

Addendum 1

The bias correction applied to these models would be useful for quantitatively associating the difference between real and corrected predicted ages with a decline and reserve in cognitive function. However, owing to the limitations of this research pertaining to the practice of screening brains (Brain Dock), the data used in this study were not based on random sampling but hinged on the selection criteria in that the participants had undergone that health checkup service. As mentioned in the main text, the main target participants were healthy senior citizens who tended after retirement to vigorously take care of themselves, while many middle-aged patients were facing health concerns triggered by the notification of results of medical examination held at their workplaces. Presumably due to the irregular difference in mindset and feeling between the elderly and worried middle-aged patients, the phenotype data were heterogeneous in the first place, showing that, quite counterintuitively, there was a negative correlation (P = 0.001) between apathy scores and actual ages: the retired elderly people were more motivated than the younger people still working. However, data modeling allowed us to amend the evaluation of objective brain ages by rectifying that unlikely direction of correlation at least weakening the effects of participants’ idiosyncrasy. The difference between actual and corrected ages and apathy scores were positively correlated for all the modeling outcomes, although the results were not significant. The apathy scores and the difference between actual and predicted ages corrected by techniques 1 and 2 were righteously correlated but with P = 0.085 and 0.079, respectively.

## Appendix 2: Gender classification

**Figure A2.**
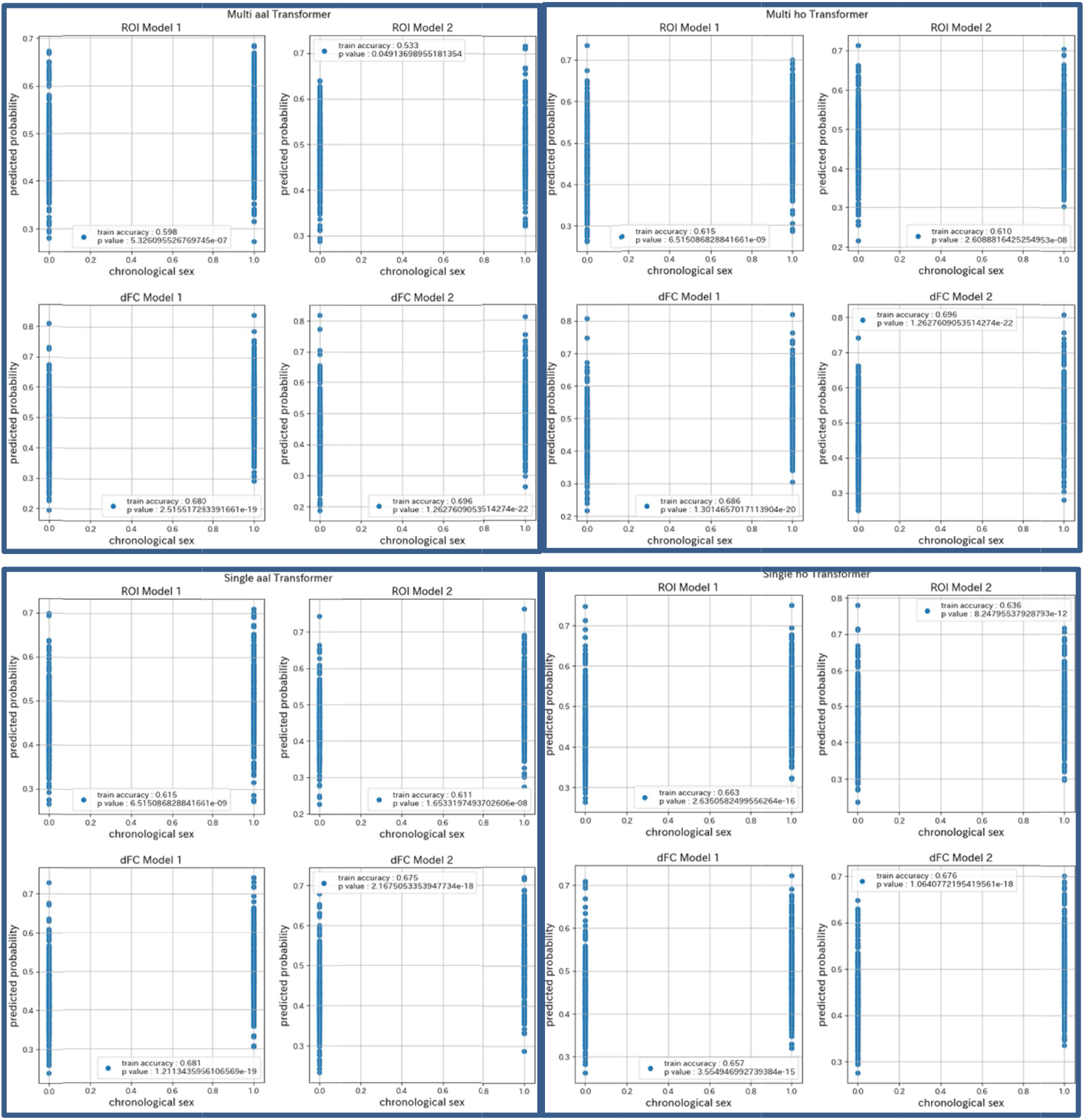
Gender discrimination results (predicted probability and training accuracy) compared between the multi-task and single-task models. Panels above: multi-task, below: single-task models.

**Table A5.** Accuracies of gender classification using the multi-task Transfo**r**mer model

| Multi-task | ROI 1<br>accuracy | ROI 2<br>accuracy | dFC 1<br>accuracy | dFC 2<br>accuracy |
| --- | --- | --- | --- | --- |
| Transformer AAL | 0.5984** | 0.5333* | 0.6797** | 0.6959** |
| Transformer HO | 0.6146** | 0.6098** | 0.6862** | 0.6959** |

\*\*P<0.01, \*P<0.05

**Table A6.** Accuracies of gender classification using the single-task Transformer model

| Single-task | ROI 1<br>accuracy | ROI 2<br>accuracy | dFC 1<br>accuracy | dFC 2<br>accuracy |
| --- | --- | --- | --- | --- |
| Transformer AAL | 0.6146** | 0.6114** | 0.6813** | 0.6748** |
| Transformer HO | 0.6634** | 0.6358** | 0.6569** | 0.6764** |

\*\*P<0.01, \*P<0.05

## Appendix 3: Codes

### Transformer algorithm

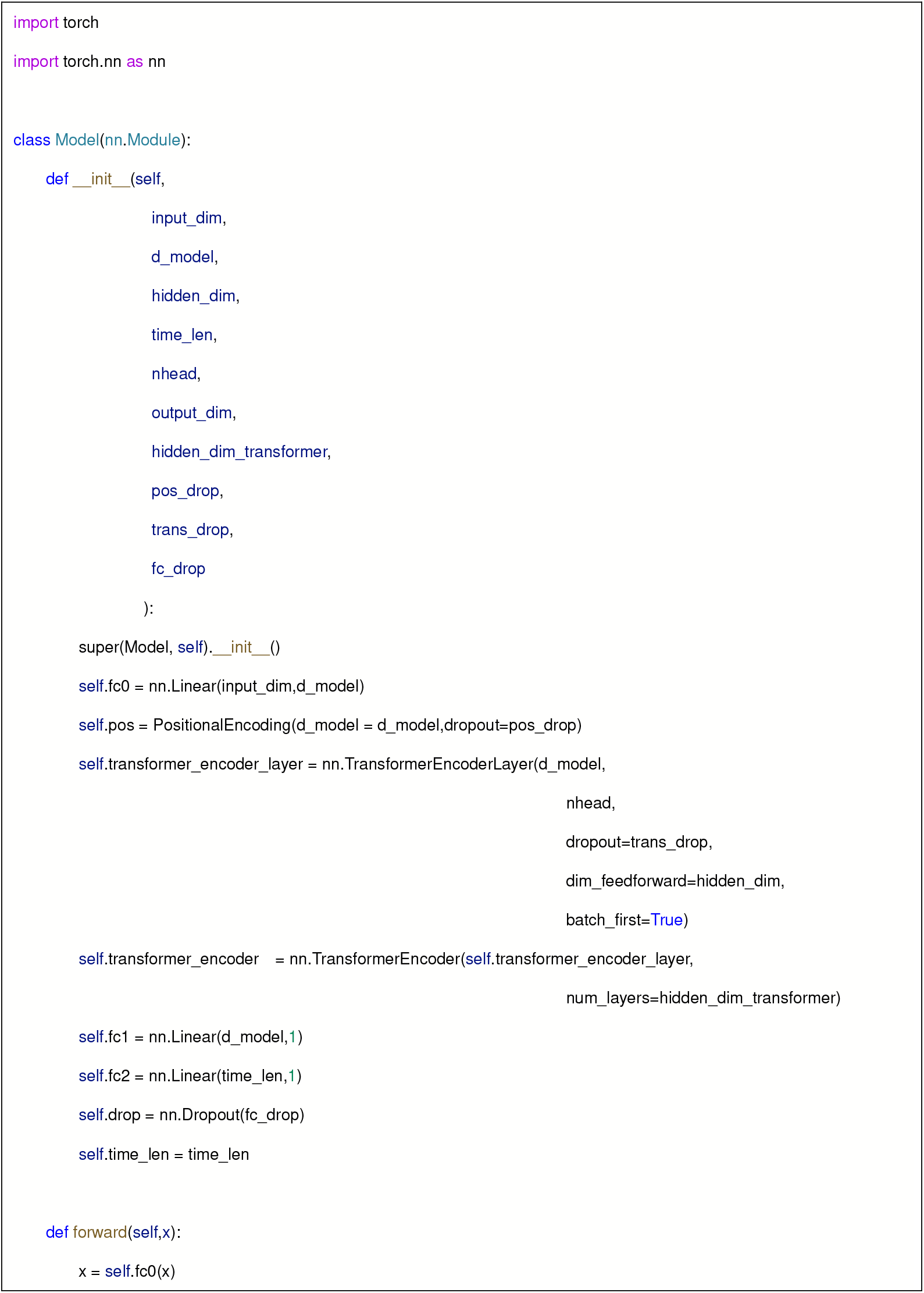
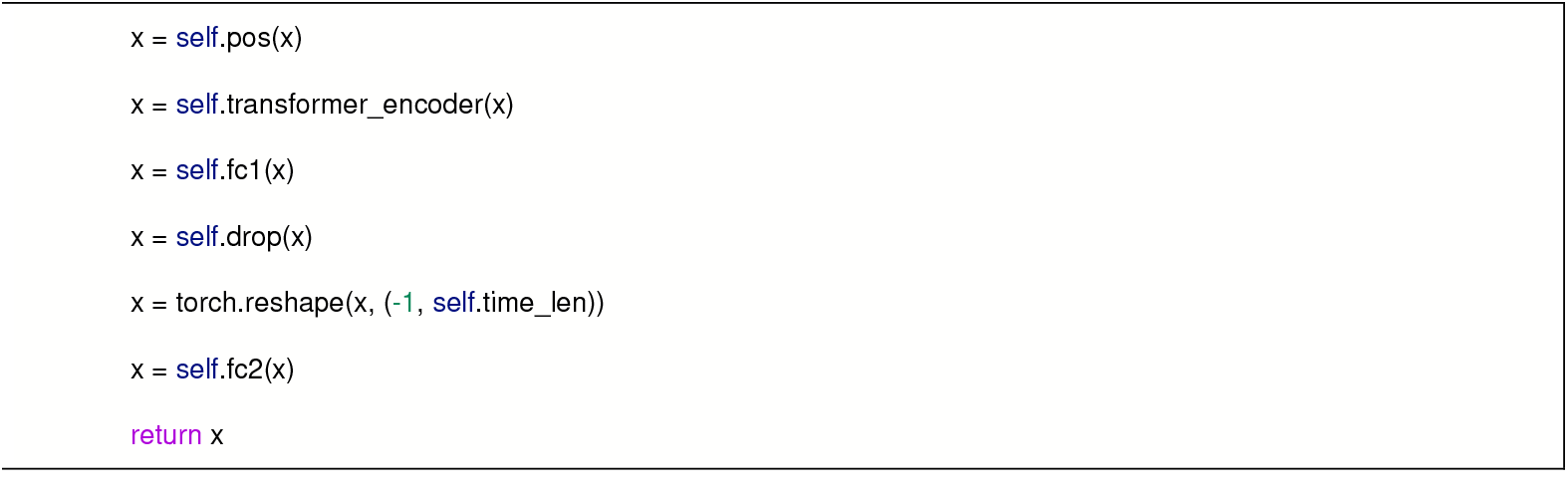

### Multi-task Transformer algorithm

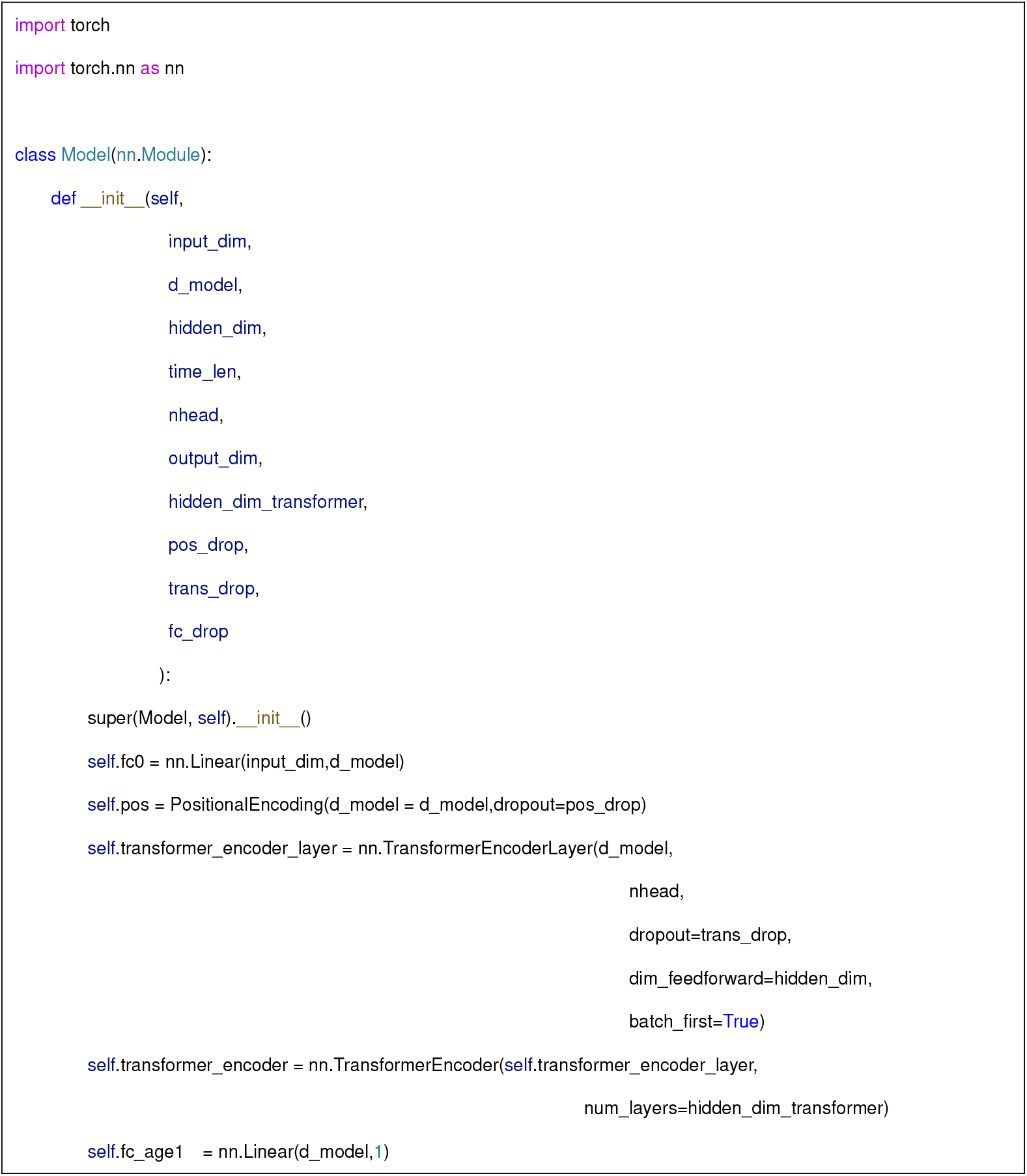
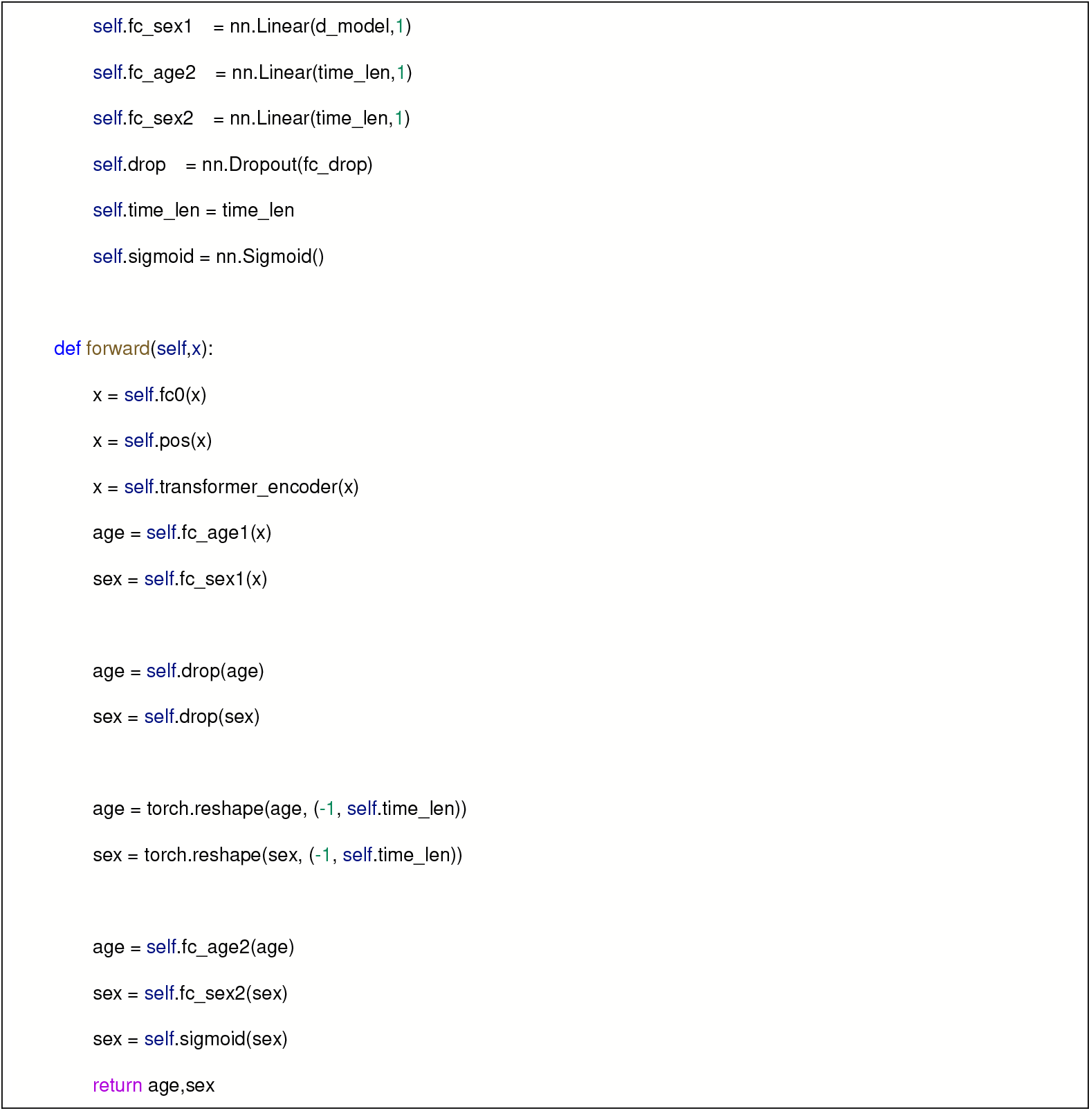

### LSTM algorithm

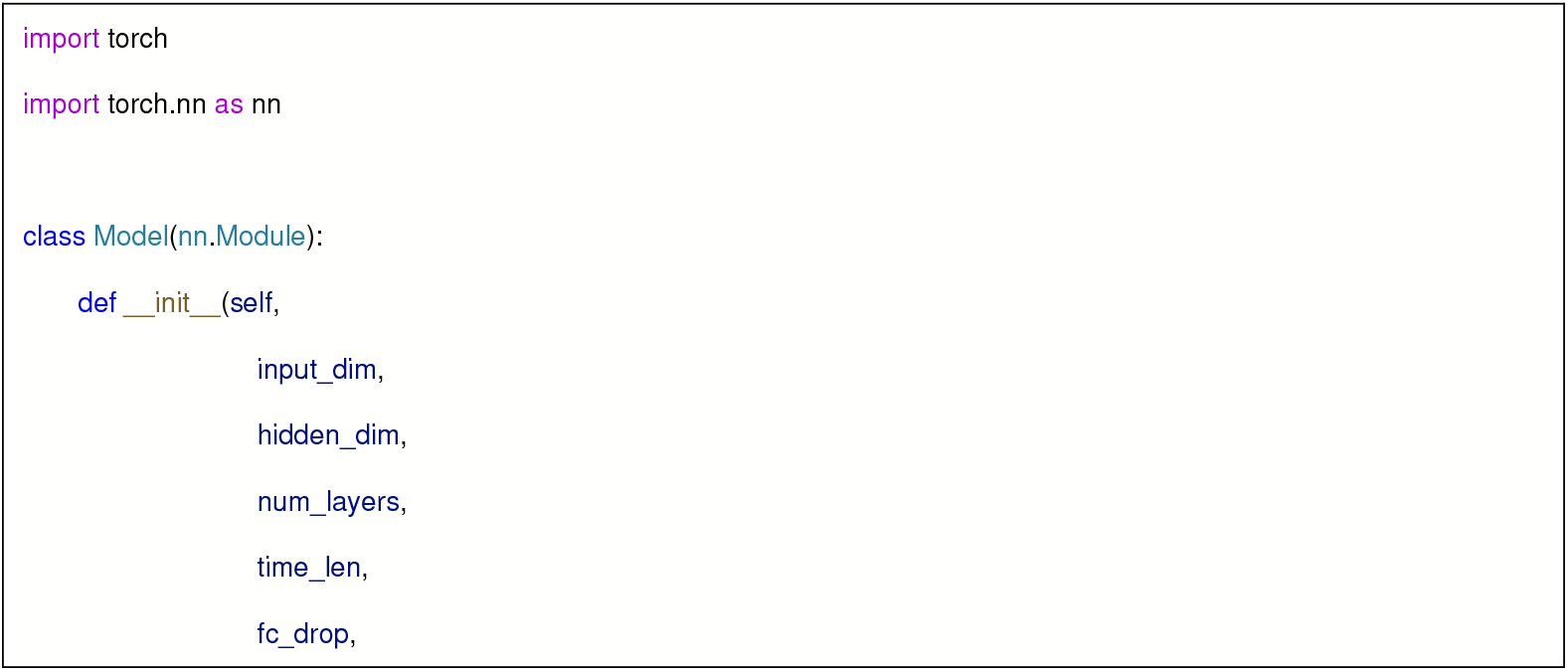
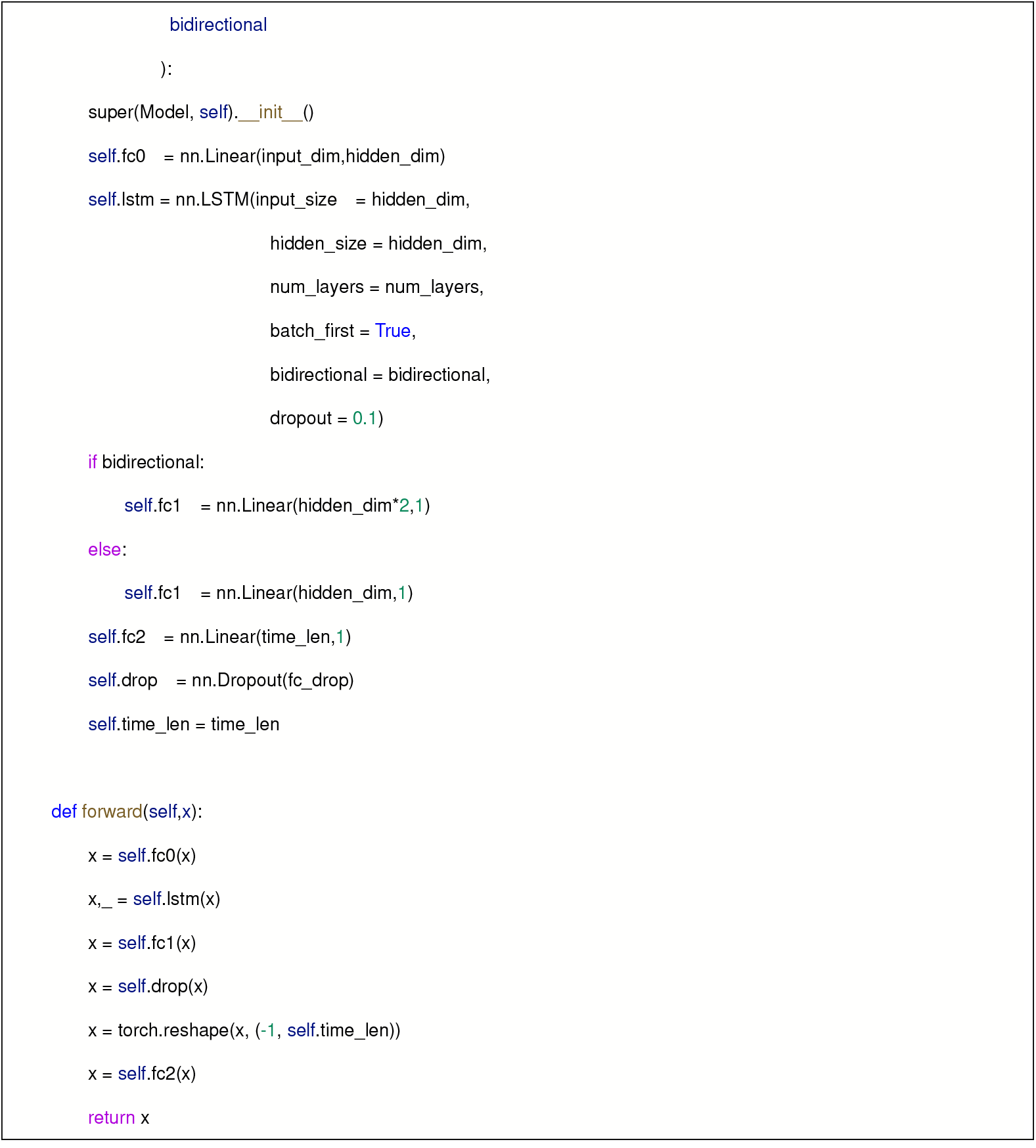

### Loss functions and learning steps of the Transformer and LSTM modeling algorithms

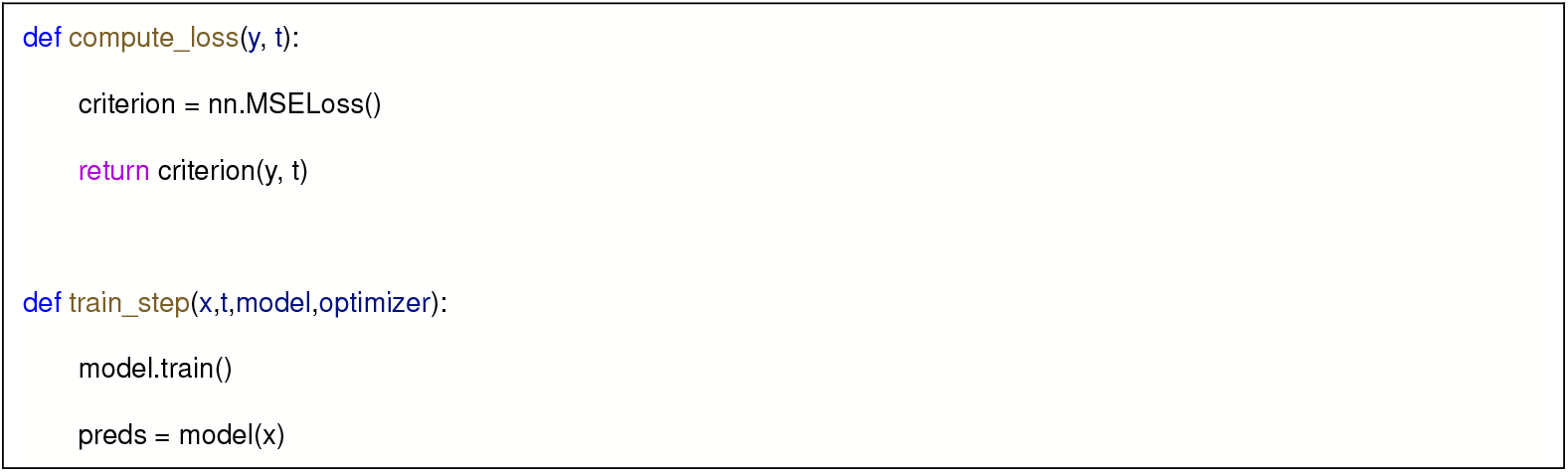
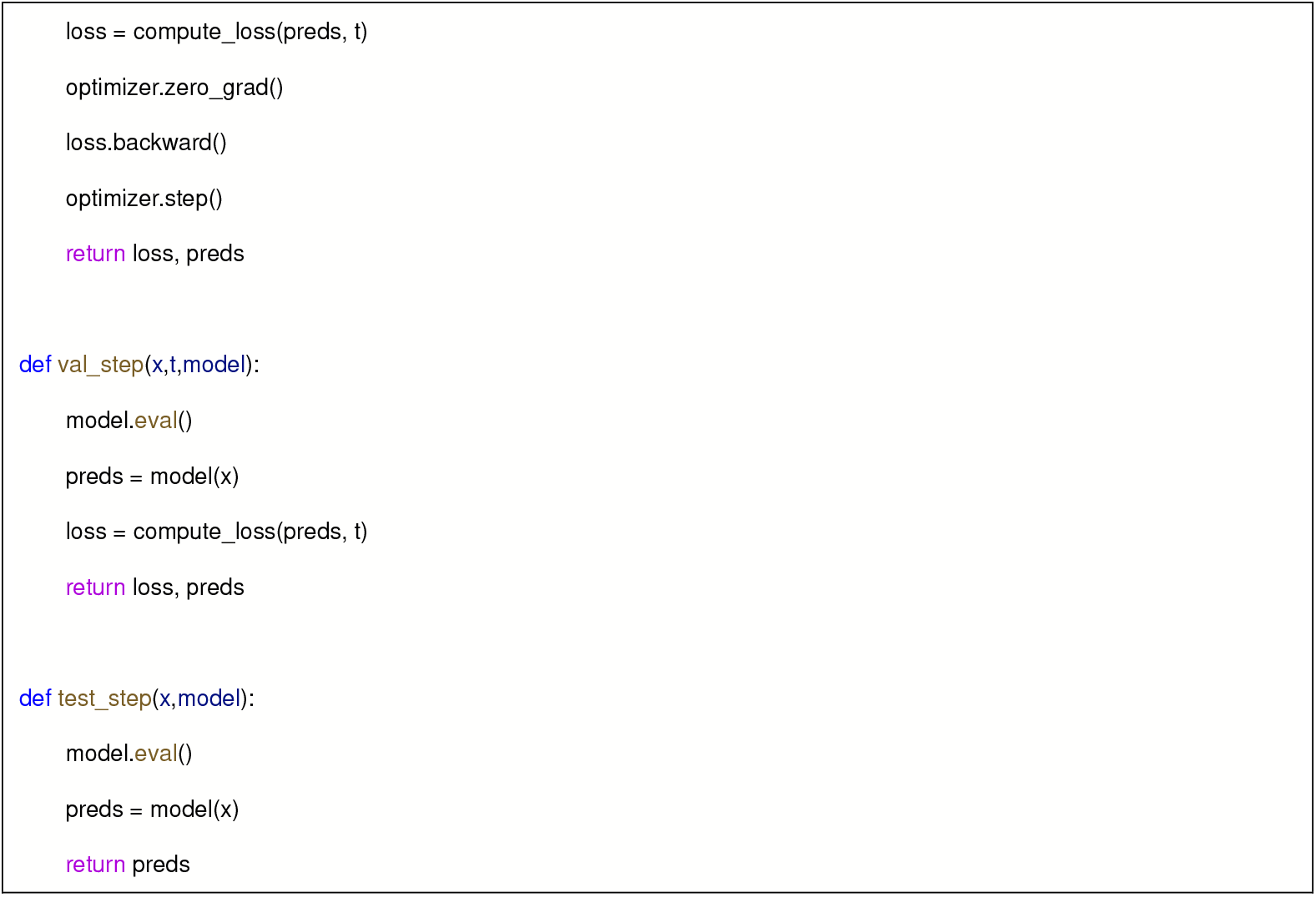

### Loss functions and learning steps of the Multi-task Transformer modeling algorithm

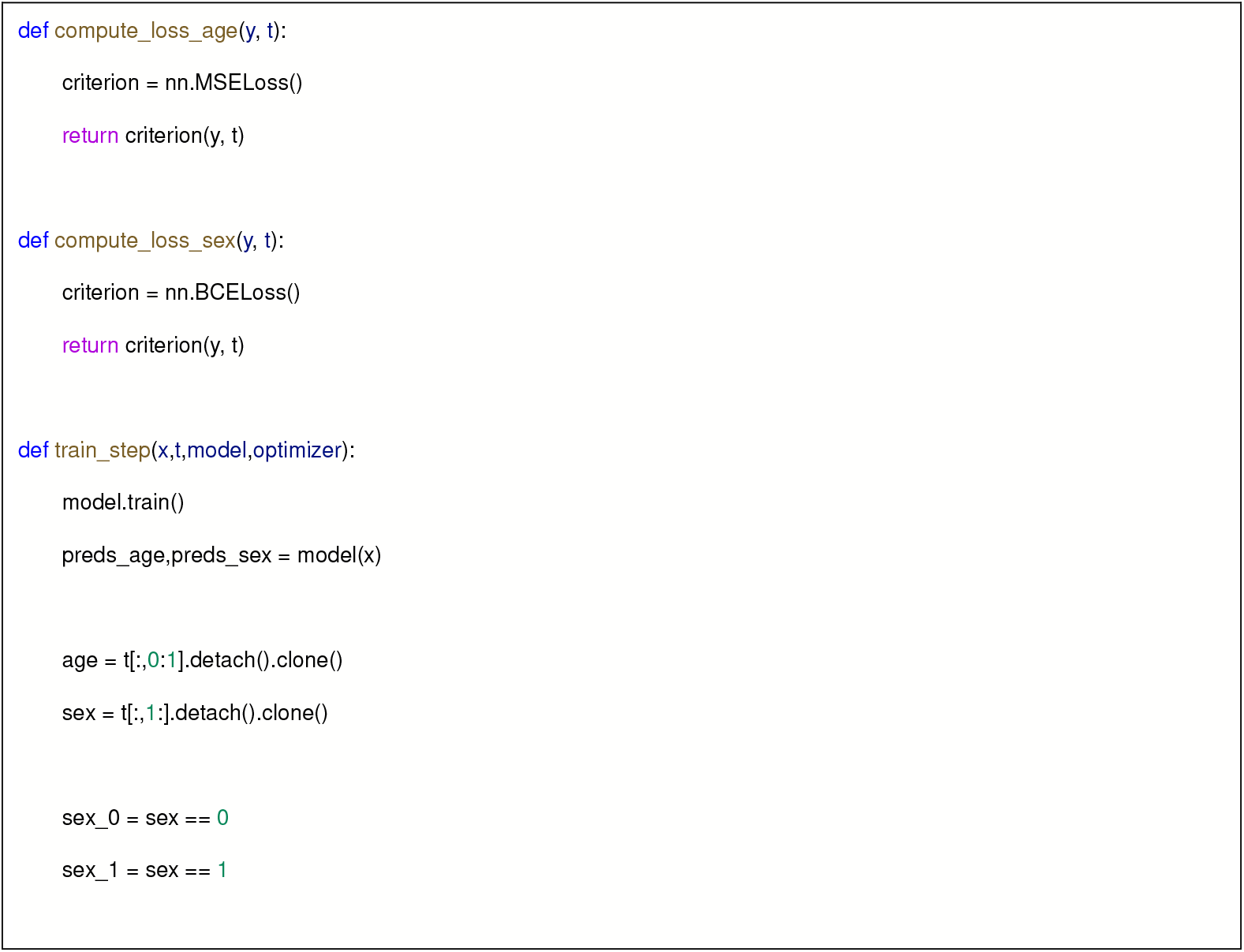
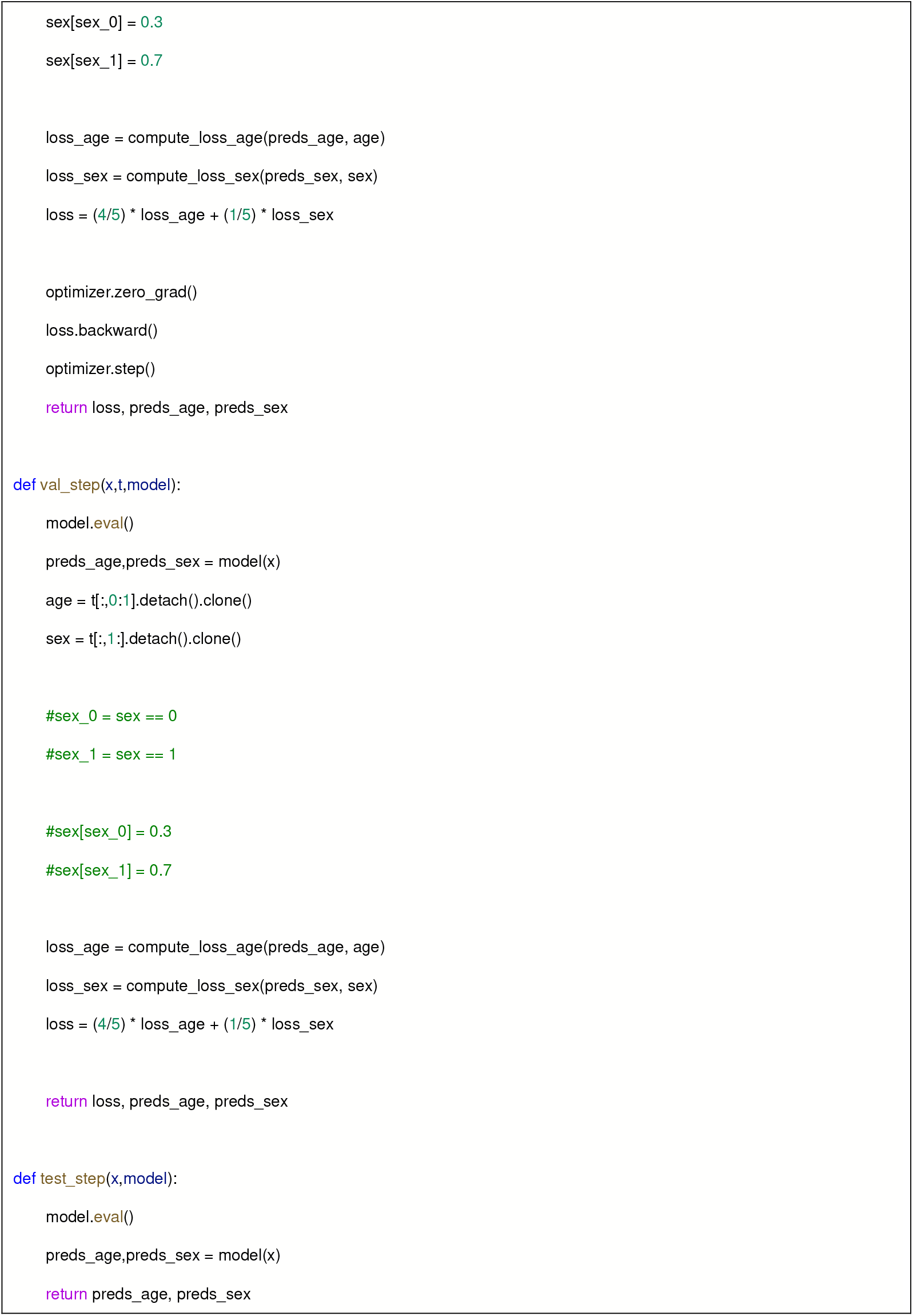

